# The Dark Side of Mentalizing: Learning Signals in the Default Network During Social Exchanges Support Cooperation and Exploitation

**DOI:** 10.1101/2023.05.03.538867

**Authors:** Timothy A. Allen, Michael N. Hallquist, Alexandre Y. Dombrovski

**Affiliations:** Department of Psychiatry, University of Pittsburgh, Pittsburgh, PA, 15213; Department of Psychology and Neuroscience, University of North Carolina at Chapel Hill, Chapel Hill, NC, 27599

**Keywords:** reciprocal cooperation, social learning, mentalizing, individual differences, default network

## Abstract

The evolution of human social cognitive capacities such as mentalizing was associated with the expansion of frontoparietal cortical networks, particularly the default network. Mentalizing supports prosocial behaviors, but recent evidence indicates it may also serve a darker side of human social behavior. Using a computational reinforcement learning model of decision-making on a social exchange task, we examined how individuals optimized their approach to social interactions based on a counterpart’s behavior and prior reputation. We found that learning signals encoded in the default network scaled with reciprocal cooperation and were stronger in individuals who were more exploitative and manipulative, but weaker in those who were more callous and less empathic. These learning signals, which help to update predictions about others’ behavior, accounted for associations between exploitativeness, callousness, and social reciprocity. Separately, we found that callousness, but not exploitativeness, was associated with a behavioral insensitivity to prior reputation effects. While the entire default network was involved in reciprocal cooperation, sensitivity to reputation was selectively related to the activity of the medial temporal subsystem. Overall, our findings suggest that the emergence of social cognitive capacities associated with the expansion of the default network likely enabled humans to not only cooperate effectively with others, but to exploit and manipulate others as well.

**Significance:** To navigate complex social lives, humans must learn from their interactions with others and adjust their own behavior accordingly. Here, we show that humans learn to predict the behavior of social counterparts by integrating reputational information with both observed and counterfactual feedback acquired during social experience. We find that superior learning during social interactions is related to empathy and compassion and associated with activity of the brain’s default network. Paradoxically however, learning signals in the default network are also associated with manipulativeness and exploitativeness, suggesting that the ability to anticipate others’ behavior can serve both the light and dark sides of human social behavior.

---

Relative to other species, humans’ social lives involve extraordinary levels of cooperation. Anthropoids began living in loose social groups to reduce predation risk over 50 million years ago, and hominins have evolved increasingly sophisticated forms of sociality even more recently (Shultz et al., 2011). This transition to sociality conferred major advantages, but also exposed individuals to intergroup rivalries (Dunbar, 2012). Selection pressures thus favored social cohesiveness, with individuals forging cooperative bonds to reduce conflict and coordinate foraging, rearing, and defense (Dunbar & Shultz, 2010). The computational demands imposed by this social environment were a key driver of the recent expansion of the human cortex, and particularly the default network (Dunbar, 2009).

An apparent paradox in the evolution of cooperative social behavior is that actors who consistently behave selfishly in social exchanges have an individual fitness advantage over unselfish conspecifics (Fletcher & Doebeli, 2008). Yet, *populations* comprised entirely of unselfish actors have higher average fitness than those comprised exclusively of selfish actors (Nowak, 2006). How can these seemingly divergent predictions at the individual and population levels be reconciled? One mechanism that maintains unselfish behavior at the population level is reciprocity. In reciprocal exchanges, actors reward kindness and punish defection or aggression, assuming that the temporary costs of prosocial actions will be offset by benefits returned in subsequent exchanges (Axelrod & Hamilton, 1981; Trivers, 1971). Game-theoretic studies show that in repeated social interactions where individuals have a choice between cooperating or defecting, reciprocity can be evolutionarily stable and resistant to incursion by other strategies (Axelrod & Hamilton, 1981). Still, no single strategy is optimal, and phenotypic heterogeneity in cooperation is common (Nowak, 2006; Zheng et al., 2017). For instance, unconditional defection is also a stable strategy (Axelrod, 1981). Altogether, whereas social cooperation generally supports the survival of human social groups, individuals acting selfishly at the expense of the group can fare remarkably well.

In humans, variability in cooperative behavior is described by a dimension of personality known as a*greeableness*. Agreeableness reflects the tendency to coordinate one’s own goals with those of others, enabling individuals to collaborate effectively while avoiding aggressive or threatening social interactions (Allen & DeYoung, 2017; Graziano & Tobin, 2009). Conversely, the low pole of agreeableness, *antagonism*, is a central feature of clinical personality disorders that predicts difficulties with attachment and intimacy, aggression and hostility toward others, and, in treatment, fractures in the therapeutic relationship (Ringwald et al., 2021; Wright et al., 2022).

Agreeableness-antagonism consists of two lower-order traits. Compassion-callousness contrasts empathy and generosity with indifference and cruelty. Politeness-exploitativeness, contrasts honesty and humility with selfishness and manipulativeness (Allen et al., 2020). A population varying on these traits can display a mix of strategies for navigating social relationships. For instance, humans engage in reciprocal cooperation both to build connection (high compassion) and to coerce social partners into behaving in ways that benefit their own fitness (high exploitativeness; Bereczkei, 2018). Conversely, a callous individual may eschew social bonds altogether and act solely to maximize individual fitness (Axelrod & Hamilton, 1981), whereas a polite person may display unconditional prosociality and kindness (Thielmann & Hilbig, 2015; Zhao et al., 2016).

Individual differences in compassion-callousness and politeness-exploitativeness describe the various strategies humans use to navigate social relationships. But how do these strategies emerge dynamically in social exchanges? The study of learning during social exchanges can uncover the computations and neural mechanisms underlying both human social cooperation and exploitation. In previous work, we found that humans learn from social exchanges by considering the counterfactual outcomes of their untaken actions (Vanyukov et al., 2019). Rather than learning from rewards alone, knowing what could have happened in an exchange allows one to update the value of one’s policy, or approach, toward a social partner (see Figure 1). This “policy model” explains why, in the interpersonal context, it is often rewarding to not only predict a counterpart’s cooperative actions but also their defections, as when one elects not to ask out an acquaintance and then later learns from a mutual friend that the acquaintance was never interested. In this instance, the policy model predicts that the decision-maker experiences a *positive* prediction error (PE) because the experienced outcome (no date) is better than it would have been had they taken an alternative action (making an overture and having it rejected). This positive PE, in turn, updates the individual’s underlying policy in future interactions with the acquaintance.

**Figure 1.**
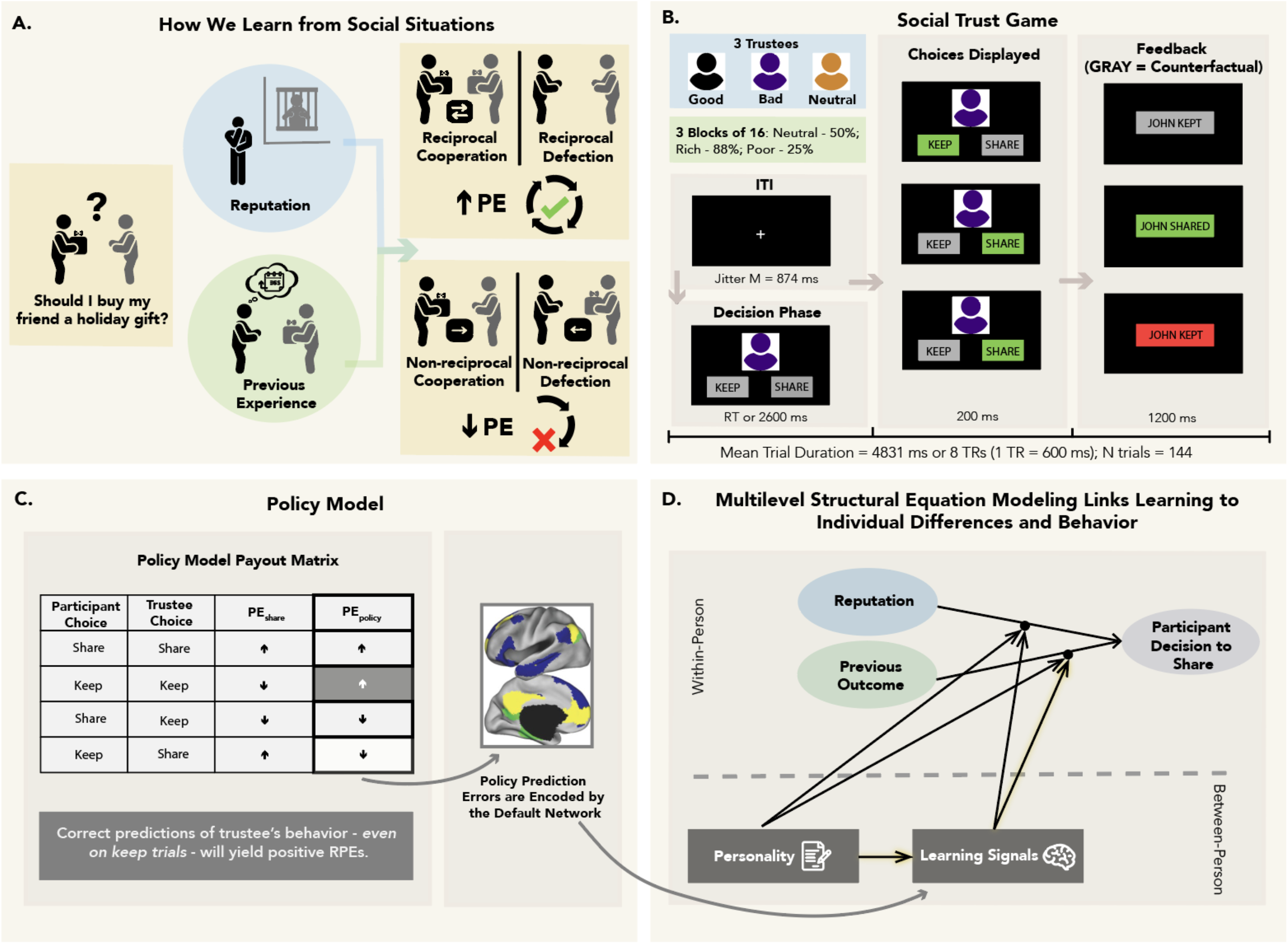
Rationale, design, and analytic approach. Individuals learn from experience by selecting an action, observing its outcome, and updating the expected reward value of future actions. Value updates are made using prediction errors (PEs), which reflect the discrepancy between expected and obtained outcomes such that better-than-expected outcomes lead to positive PEs and worse-than-expected outcomes lead to negative PEs. Other salient social information can also be integrated with experience to influence social decision-making, as when the reputation of one’s partner predicts decisions to trust them even when reputation is unrelated to the partner’s actual behavior (Fouragnan et al., 2013; Vanyukov et al., 2019). A) Consider the decision about whether to buy a friend a holiday gift. Reputational information (e.g., news of a friends’ immoral behavior; blue circle) can be integrated with reinforcement history (e.g., did the friend buy you a gift last year? green circle) to affect one’s policy toward their social counterpart. Critically, decisions that correctly anticipate the behavior of one’s social partner (correctly predicting they bought you a gift, or correctly predicting they *did not* buy you a gift) yield positive reward prediction errors, leading to a cycle of reciprocity even when no actual reward is received. B) Participants played a modified iterative social trust game with three fictional trustees, in which they had the option to keep an initial endowment or share it in the hopes of increasing their profit if the trustee also shared. Pre-task vignettes were used to manipulate trustees’ reputations (blue box). To manipulate reinforcement history, trustees shared at varying rates across rich, poor, and neutral blocks (green box). On trials where participants kept, counterfactual feedback about what the trustee would have chosen was provided, even though it did not affect the trial payout. C) The policy model posits that individuals learn from social feedback by using counterfactual reasoning to optimize their approach, or policy, toward their social counterpart, leading to better anticipation of the counterpart’s behavior. One implication of this is that correct predictions of the trustee’s behavior will lead to positive reward prediction errors, even if no actual reward is provided. This can be seen by comparing the expected direction of PEs in a model in which participants track actual rewards (column 3) vs. a model in which they track the success of their policy toward the counterpart (column 4). We propose that policy prediction errors are primarily encoded within the brain’s default network. D) Multi-level structural equation modeling (MSEM) was used to evaluate whether between-person variables (e.g., policy prediction errors encoded in the default network) moderate the effect of design variables on trial-level decision making. Personality traits were introduced as between-person predictors of policy PEs in the default network. Formal tests of mediation were then used to examine the indirect effect of traits on behavior via learning signals (highlighted lines). Default network figure adapted from Andrews-Hanna et al., 2014. Note: trustee faces were taken from the NimStim Face Stimulus Set (Tottenham et al., 2009) for the experiment, but were changed to silhouettes here at the request of the preprint server.

Counterfactual learning, which improves predictions of a social counterpart’s behavior, facilitates mentalizing (Byrne, 2016). Mentalizing, in turn, is associated with greater reciprocal cooperation (Barrett et al., 2010). For instance, in a social exchange, agents reason about the intentions of others before reciprocating (Falk & Fischbacher, 2006; McCabe et al., 2003; Stanca et al., 2009; van den Bos et al., 2011). Interestingly, performance on mentalizing or theory of mind tasks is negatively associated with callousness, but positively associated with exploitativeness (Allen et al., 2017). This dissociation may reflect differences in counterfactual learning in the social setting. Specifically, individuals low in callousness and high in exploitativeness may rely on counterfactual thinking to anticipate how a counterpart will behave, adjusting their own decisions accordingly. This reasoning informs our hypothesis that callousness is negatively, and exploitativeness positively, associated with sensitivity to PEs during social exchanges, leading to better anticipation of a counterpart’s decisions.

A key neural substrate of human mentalizing and cooperation is the default network, a large-scale functional network that spans the medial prefrontal cortex, posterior cingulate cortex, inferior parietal lobule, and medial and lateral regions of the temporal cortex (Buckner et al., 2008). Default network activity has been linked to higher agreeableness (less antagonism) and better performance on mentalizing tasks (Udochi et al., 2022). Genetic and comparative neuroimaging studies have shown that the default network expanded disproportionately in recent human evolution (Buckner & Krienen, 2013; Hill et al., 2010), in part due to the expression of genes that diverge with the human lineage and are associated with sociability (Wei et al., 2019). Consistent with this, the structure and connectivity of the default network scales with social network size (Noonan et al., 2018).

The default network supports social cognition and self-generated thought, including episodic memory, mental simulation, and mentalizing (Andrews-Hanna et al., 2014). It is also activated by episodic counterfactual thinking, or simulating what could have happened if one had acted differently in the past (van Hoeck et al., 2015). Indeed, we have previously found policy PE signals during social exchanges – which signify surprise about the counterpart’s behavior relative to one’s prediction – in default network regions including the posterior cingulate and precuneus (Vanyukov et al., 2019).

Using a computational model of learning during social exchanges, we investigated whether learning signals encoded by the default network that update the value of one’s policy toward a social partner are associated with individual differences in callousness and exploitativeness and their behavioral expression. Participants, including individuals diagnosed with borderline personality disorder, completed a modified iterative social trust game in which they interacted with trustees of varying reputations. Integrating task-based fMRI, cognitive computational modeling, and multilevel structural equation modeling, we show that learning in the default network is negatively related to callousness and positively related to exploitativeness. Moreover, heterogeneity in default network activity accounts for the two facets’ distinct effects on reciprocity during social exchanges.

## Results

### Modified Trust Game: Behavior

In a modified social trust game (Figure 1B), participants interacted with three different trustees of varying reputation (good, bad, neutral; 48 trials each, 144 trials total). On each trial, participants received $1.00 to keep or share with the trustee. If they chose to keep, they retained their dollar. If they chose to share, the outcome was contingent on the trustee: if the trustee also shared, they received $1.50, whereas if the trustee kept, they were left with nothing. All trustees shared 50% of the time in the first block of 16 trials, and then either 25% (poor block) or 88% (rich block) of the time in two subsequent counterbalanced blocks of 16 trials. Critically, to enable participants to learn from counterfactual outcomes, they were shown the trustee’s decision even after choosing to keep the $1.00.

Participants were less likely to share with the bad trustee than neutral or good trustees (neutral: *β* = -.24, *SE* = .05, *p* < .001; good: *β* = -.26, *SE* = .05, *p* < .001). There was also a reciprocity effect (*β* = .62, *SE* = .07, *p* < .001) such that when trustees shared on one trial, participants often shared on the next, whereas when trustees kept the investment, participants often kept on the subsequent trial (full behavioral results in Supplement Section IV). Because the rate of cooperation did not vary between the good and neutral trustees, these conditions were collapsed into a single category for subsequent analyses.

### Computational Model of Learning During Social Exchanges

We used computational modeling to examine whether, in a social interaction, participants learn by considering the counterfactual outcomes of their untaken actions in addition to the actual rewards they receive in each trial (as proposed by the policy model; see Figure 1C). Additionally, we were interested in whether the policy model could capture individual differences in the overall rate of cooperation and in participants’ sensitivity to the reputation of their social partner. We tested whether the policy model provided a better fit to behavior on the task compared to three competing models. All models included the same five subject-specific parameters: 𝛼, a learning rate parameter reflecting the speed at which participants update the value of keeping or sharing; *b*, a temperature parameter reflecting choice stochasticity, *k_s_*, a participant-level parameter reflecting a bias of the participant to keep or share regardless of reinforcement; and two condition-level *k_t_* parameters, reflecting the participant’s bias to keep or share with the good or bad trustee relative to the neutral trustee. Critically, the models differed in terms of how reinforcement on the task was encoded (i.e., each model had a different payoff matrix; see Supplement Section VI). The key prediction of the policy model was that participants would learn not only from actual rewards on each trial, but also from counterfactual feedback provided on keep trials. Bayesian model comparison of the four competing models indicated that the policy model dominated all alternatives (BOR < .001; protected exceedance probability for the policy model = 1.00), replicating our previous findings (Vanyukov et al., 2019). Parameters from the winning policy model were minimally intercorrelated (Figure S4A).

Next, we examined whether participant-level parameters derived from the policy model related to individual differences in antagonism and its two facets, callousness and exploitativeness (traits were derived using exploratory factor analysis, see Figure 3A & Supplement Section V for details). Antagonism was negatively related to the learning rate (*β* = -.05, *SE* = .02, *p* = .02) and the participant-specific *κ_s_* parameter (*r* = -.21, *p* = .01), though the latter effect was attenuated when controlling for age (*β* = -.07, *SE* = .04, *p* = .054). This is consistent with antagonistic individuals being slower to update keep/share values based on new information and less likely to share in general (Figure S4A-B). At the facet level, exploitativeness (controlling for callousness) was negatively related to the *κ_s_* parameter (*r* = -.19, *p* = .02), though again, this effect did not withstand correction for age (*β* = -.05, *SE* = .05, *p* = .31). Callousness (controlling for exploitativeness) was positively associated with the *κ_t_* parameter for the bad trustee (*β =* .05, *SE =* .02, *p* = .03), suggesting a bias toward sharing with the bad trustee (Figure S4C). This finding mirrored model-free behavioral results in which callousness, but not exploitativeness, moderated the effect of reputation on sharing (*β* = .26, 95% CI = [.04, .45], *p* = .01; Table S12). Participants who were low (−1 *SD*) or average on callousness were less likely to share with the bad trustee relative to the good and neutral trustees, whereas reputation had no effect on sharing for those high (+1 *SD*) in callousness (Figure S7C).

**Figure 3.**
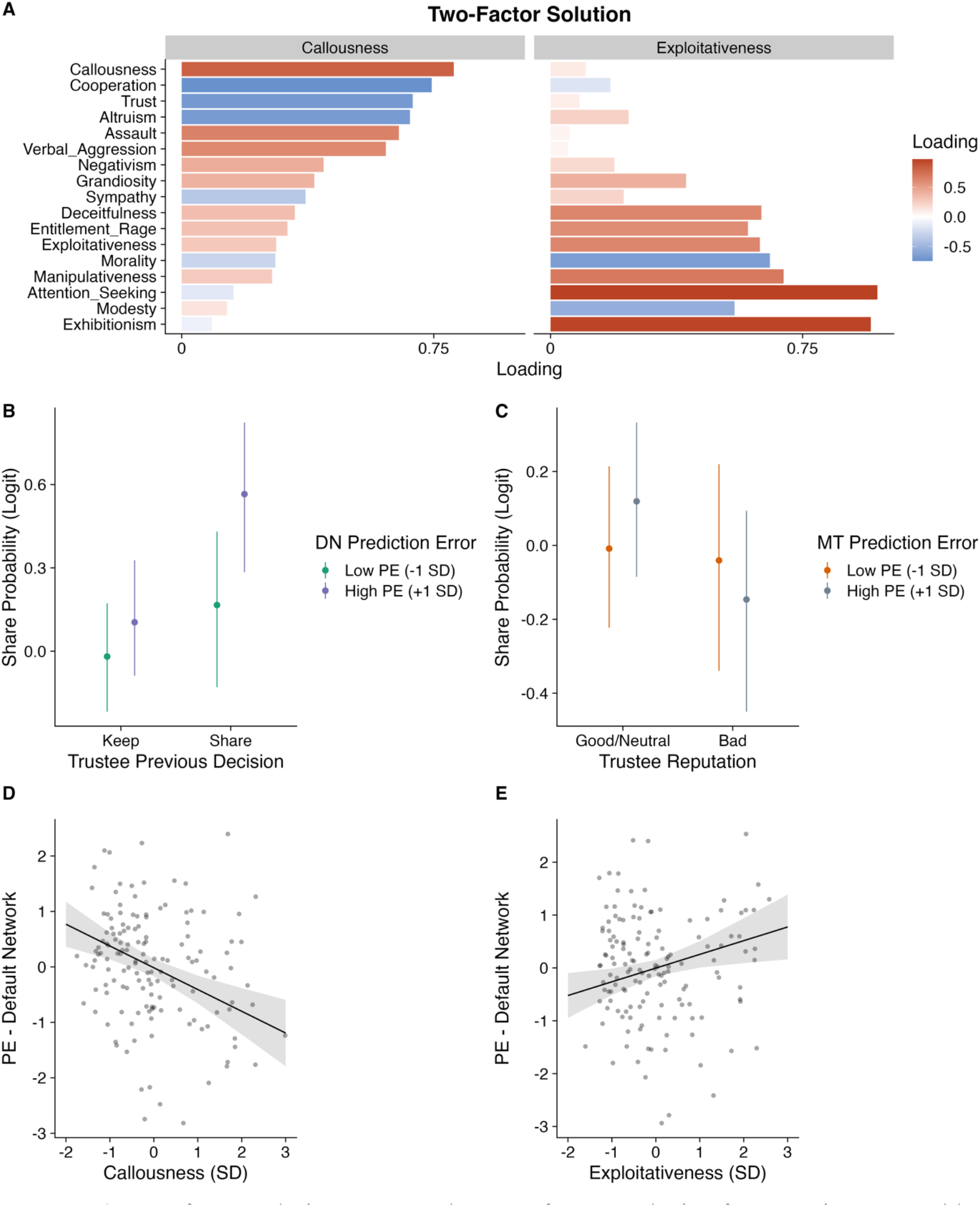
A) Two-factor solution to an exploratory factor analysis of antagonism-agreeableness scales. B) Policy PEs in the default network moderate the effect of the trustee’s last decision on the participant’s choice to keep or share. C) Policy PEs in the medial temporal subsystem moderate the effect of trustee reputation on participant decisions. D) Callousness was negatively, and E) exploitativeness was positively, associated with policy PEs in the overall default network. Abbreviations: PE = prediction error; DN = default network; MT = medial temporal subsystem; SD = standard deviation.

### Policy Prediction Error Signals in Default Network

Based on our previous results and studies implicating the default network in counterfactual reasoning and mentalization, we expected to find policy PE signals both in the canonical striato-limbic reward prediction error network and in the default network. Fig 2A depicts the whole brain prediction error map (FWE *p* < .05). Signed PEs were found bilaterally in the ventral and dorsal striatum, putamen, as well as in the lateral occipital cortex, pre- and postcentral gyrus, cerebellum, and the frontal operculum. PEs were also evident in regions comprising the default network, and especially the posterior hub, where we observed significant PE activation in 85% of voxels. PEs were also present in the anterior cingulate cortex, precuneus, and bilateral middle and inferior temporal gyrus (Supplemental Table S6 provides a complete listing of surviving clusters). Overlap between PEs from the policy model and the canonical default network is depicted in Fig 2B.

**Figure 2.**
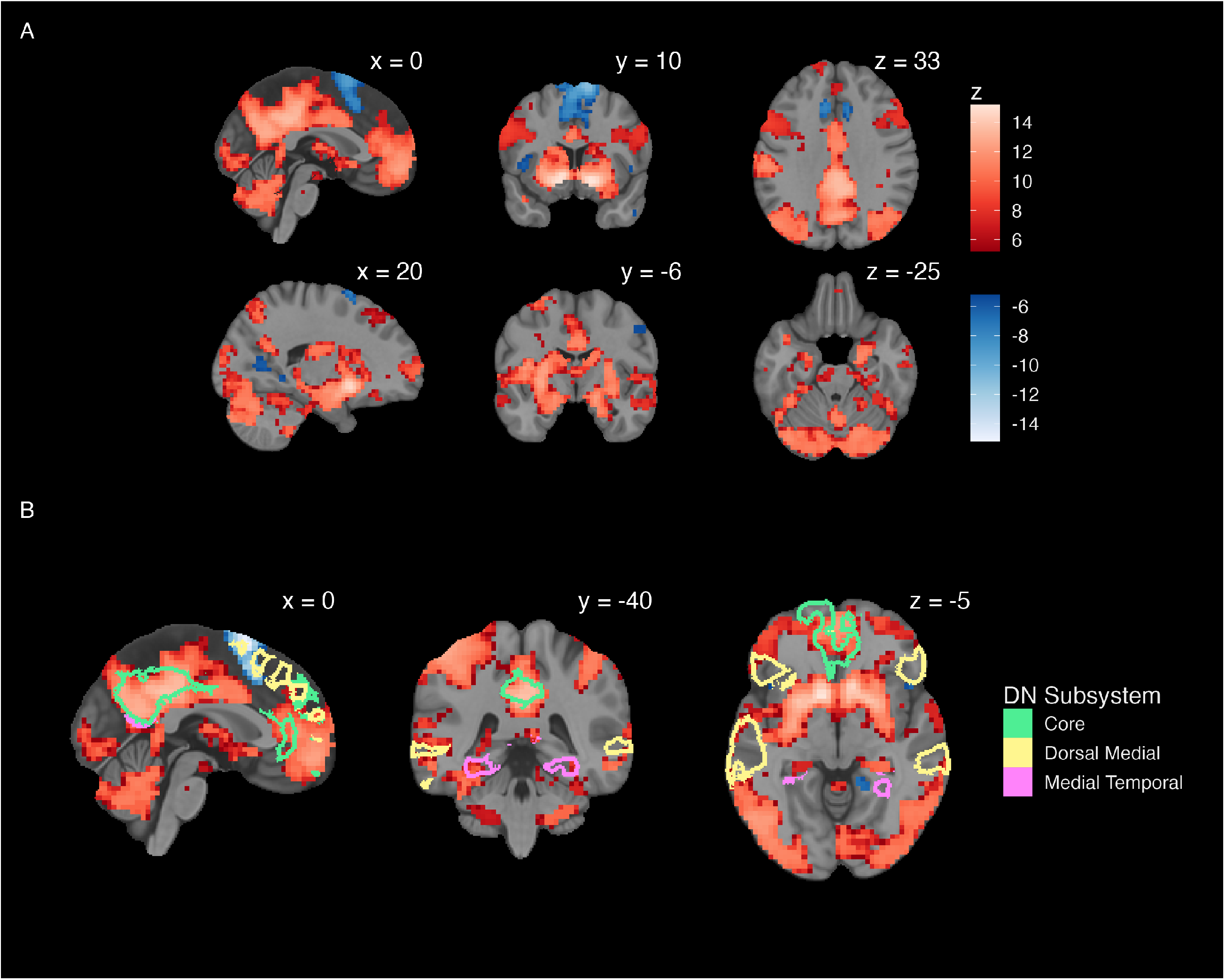
A) Whole brain prediction error map. Threshold z > 5.17, p_ptfce_ < .05. B) Overlap between surviving prediction error clusters and regions of the canonical default network, broken down by subsystem. Images created using *ggbrain* (Hallquist, 2022b).

### Policy Prediction Error Signals in the Default Network and Reciprocity

Since learning from counterfactual feedback facilitates reciprocal cooperation, we expected policy PEs would be associated with greater reciprocity in social exchanges. To test this prediction, we fit a multilevel structural equation model (MSEM) with random slopes in which individual differences in the strength of PE modulation in the default network predicted the effects of exchange (trial within trustee), reputation, and trustee’s previous decision on participant decisions to keep or share (coded 0/1, respectively; Table S7). Individuals with stronger PE signals in the default network exhibited greater reciprocity (*β* = .25, 95% CI = [.09, .39], *p* = < .001). Specifically, for those with high, but not low, PE modulation of the default network, the trustee’s decision to share on the previous trial was associated with a higher likelihood of the participant reciprocating (see Figure 3B). PE modulation of the default network had no effect on the association between reputation and the participant’s decision to share.

The default network is functionally heterogenous, spanning three subsystems: core, medial temporal, and dorsal medial (Andrews-Hanna et al., 2014). As a test of anatomical specificity, we examined whether the effect of PE signaling on behavior was driven by a particular subsystem. A follow-up MSEM indicated that the effect of reciprocity on sharing was not moderated by PE signals within any single subsystem (Table S8), suggesting that the effect of learning on behavior is enacted at a network level. Notably, however, the effect of reputation on participants’ decisions to keep or share was moderated by the medial temporal subsystem, even after controlling for the core and dorsal medial subsystems (*β* = -.31, 95% CI = [-.53, -.03], *p* = .04). Specifically, the likelihood of sharing was lower with a bad (vs. good, neutral) trustee only for those with stronger PE signals in the medial temporal subsystem (Figure 3C).

### Antagonism, Callousness, Exploitativeness and Prediction Errors in the Default Network

If policy PEs enable individuals to learn about and correctly anticipate the behavioral strategies of others, they should also be related to traits linked to mentalizing, including compassion (the opposite pole of callousness) and exploitativeness. Indeed, controlling for age and sex, we found that callousness was negatively (*β* = -.39, *SE* = .10, *p* < .001) associated with prediction error modulation of the default network, whereas the reverse was true for exploitativeness (*β* = .26, *SE* = .10, *p* = .01; Figure 3D-E & Table S9). Conversely, higher-order antagonism was unrelated to prediction errors (*β* = -.10, *SE* = .08, *p* = .22). These effects also held across all three subsystems of the default network (Figure S6; the effect of exploitativeness on prediction error signaling in the dorsal medial subsystem is marginal but qualitatively similar, *p* = .05). Overall, this pattern of results suggests that antagonism’s two facets differentially predict PE modulation of the overall default network, and this effect is unlikely to be driven by a subsystem.

### Effects of Callousness and Exploitativeness on Behavior via Policy Prediction Error Signaling in the Default Network

The differential associations between the facets of antagonism and PE modulation in the default network implicate learning as a possible neural mechanism related both to self-reported personality and reciprocal decision-making during social exchanges. To formally test this, we extended the MSEM reported above by allowing the two facets of antagonism to predict PE modulation of the default network and testing whether prediction error signaling *mediated* the effect of callousness and/or exploitativeness on reciprocity. There was a significant indirect effect of both callousness and exploitativeness on reciprocity via PE signaling (see Figure 4 and Table S10 for full model results; callousness: *b* = -.05, 95% CI = -.10, -.02, *p* < .001; exploitativeness: *b* = .03, 95% CI = .004, .07, *p* = .03). These results indicate that blunted learning signals within the default network account for the association between callousness and lower reciprocal cooperation during social exchanges. In contrast, amplified learning signals in the default network link exploitativeness to greater reciprocity. Notably, these effects are only seen at the facet level, as PE modulation of the default network did not mediate the effect of higher-order antagonism on behavior (See Supplemental Section XIV).

**Figure 4.**
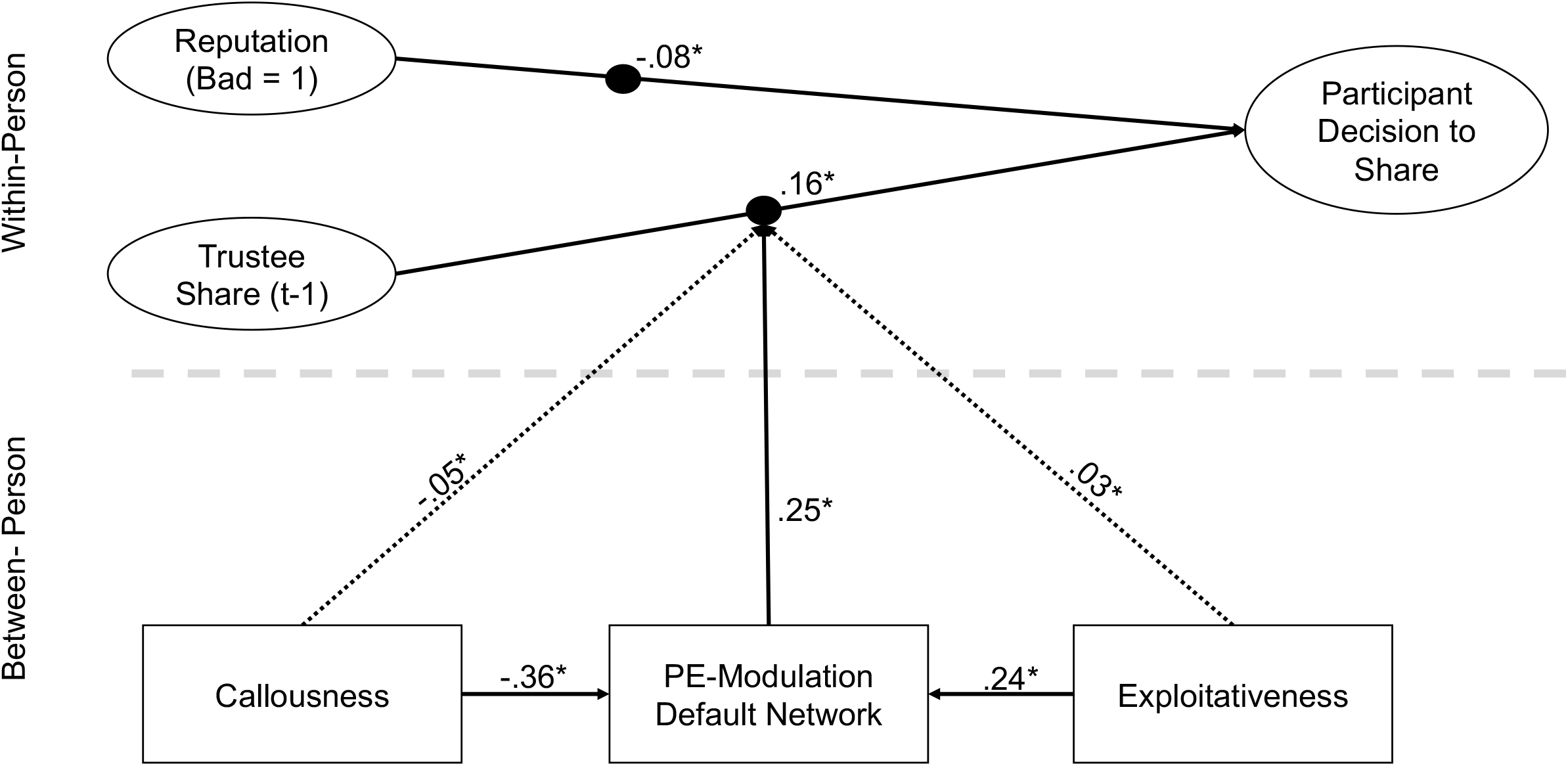
Multilevel structural equation model depicting the effect of callousness and exploitativeness on reciprocity via prediction error modulation of the default network. Boxes reflect observed variables. Circles reflect latent variables. Filled black circles indicate random slopes. Dotted lines depict indirect effects. Only significant paths are shown. Exchange number (i.e., trial within trustee, a within-person predictor), age, and sex covariates are excluded for parsimony. All coefficients are standardized except the indirect effects. * *p* < .05.

## Discussion

The evolution of human cooperation has been linked with the expansion of frontoparietal networks, and particularly the default network (Wei et al., 2019). Here, we show that cognitive capacities that emerged during this expansion and are typically thought to support prosociality can also serve the dark side of human nature, enabling manipulation, deception, and exploitation. Using a computational reinforcement learning model, we observed humans using counterfactual reasoning to learn from social feedback and predict a counterpart’s choices during an iterative trust game. More robust learning signals in the default network were seen in people who were compassionate and empathic and, paradoxically, in people who were manipulative and exploitative. These learning signals accounted for the association between each trait and reciprocity in a social exchange, consistent with the idea that humans reciprocate both to build connection and to coerce others into behaving in desired ways.

In the case of those high on compassion, superior learning from social experience facilitates cooperation and coordination, ultimately resulting in closer and more interdependent social connections. In those with high levels of exploitativeness, this same ability may be used to strategically influence others, eliciting cooperation to serve one’s immediate self-interest and preserve opportunities for long-term exploitation. Thus, while compassion and exploitativeness share an underlying capacity for learning from social experience, they are distinguished by distinct psychological functions (i.e., social connection v. personal gain). It is noteworthy that exploitativeness is associated with superior learning, as pathological traits are typically associated with learning deficits rather than advantages. However, the ability to strategically manipulate, persuade, or exploit others can be associated with greater achievement and satisfaction in certain contexts (Spurk et al., 2016). Whether superior learning and social cognition facilitate goal attainment or undermine social and emotional health in those high on exploitativeness may be highly contingent on the setting in which these capacities are employed.

Both callousness and exploitativeness were related to reciprocity on the trust task via PE modulation of the default network, but only callousness dampened the effect of reputation. It is likely that callousness reflects a fundamental insensitivity to social signals and cues, and not merely poor mentalization. Indeed, callousness is associated with behavioral deficits in recognizing others’ distress and with blunted electrocortical responses during encoding of emotional faces (Brislin et al., 2018; Brislin & Patrick, 2019). Contemporary personality theory argues that compassion-callousness fosters attachment and emotional investment in others, likely evolving as part of subcortical circuitry that facilitates interspecies pair bonding (Allen & DeYoung, 2017; Decety et al., 2016). This circuitry predates the expansion of frontoparietal networks and lateral temporal regions that accompanied more complex human mentalizing abilities.

Interestingly, we found that PE signaling in the medial temporal subsystem of the default network was associated with a stronger effect of reputation on participants’ decisions. Medial temporal regions are among the areas that experienced the least evolutionary expansion (Hill et al., 2010). Callousness was also associated with less robust learning signals in the medial temporal subsystem, and there was a marginal indirect effect from callousness to reputation via medial temporal learning signals (Table S11). The medial temporal subsystem is involved in retrieving contextual information that can inform one’s mental representation of spatial, temporal, and causal relations (Ranganath & Ritchey, 2012). By extension, our results provide tentative evidence that dampened learning signals in the medial temporal subsystem among more callous individuals represent a basic deficit in their ability to integrate relevant contextual cues, like reputation, into social decision-making.

Whereas previous studies have described functionally distinct subsystems within the default network, our results suggest that learning signals that update the value of one’s approach toward a social counterpart were encoded throughout the entire network. Indeed, no individual subsystem uniquely accounted for the default network’s effect on behavioral reciprocity or its associations with callousness and exploitativeness. Complex social dilemmas, like the one posed in our task, are likely to simultaneously recruit swaths of circuitry that integrate across subsystems and contribute to diverse cognitive processes, from episodic memory retrieval and encoding, to mental simulation, counterfactual reasoning, and mentalizing.

The posterior hub of the default network may be critical to facilitating this integration as learning occurs during social exchanges. Inspection of the overlap between our whole-brain policy PE map and the canonical default network highlighted robust modulation of the posterior cingulate cortex (PCC) and adjacent regions, including retrosplenial cortex and precuneus (85% of the voxels in the posterior hub of the default network were statistically significant). Dorsal PCC, in particular, is involved in monitoring environmental inputs to determine the need for a shift in behavioral strategy (Barack et al., 2017) and is linked with frontoparietal networks associated with goal-directed activity (Fan et al., 2019; Spreng et al., 2010).

Critically, PCC also receives projections from medial temporal regions (including hippocampus) that contribute to counterfactual scene construction (van Hoeck et al., 2015) and from components of the dorsal medial subsystem that have been linked to representing others’ mental states, including the superior temporal lobe (STGS) and temporoparietal junction (TPJ; Andrews-Hanna et al., 2014). We have found previously that connectivity between the posterior hub and STGS/TPJ is positively associated with mentalizing. Exploitativeness is also positively related to connectivity between STGS/TPJ and dorsal regions of the posterior core, whereas callousness is negatively related to STGS/TPJ connectivity with ventral regions of the posterior core (Allen et al., 2017). Taken together, our results are consistent with PCC playing a central role in integrating incoming sensory inputs from the medial temporal and dorsal medial subsystems, encoding expectancy violations (i.e., prediction errors), and flexibly adapting one’s behavioral policy via outputs to frontal cortex. From an individual differences perspective, exploitativeness and compassion may reflect a stronger influence of mentalizing information encoded in STGS/TPJ on internal representations of the environment in PCC, leading PCC to adjust behavior in the direction of greater reciprocal cooperation.

Strengths of this study include modeling trial-level behavior and decision-making using a previously validated reinforcement learning model, a well-characterized clinical sample that allowed for comprehensive psychometric assessment of the latent agreeableness-antagonism dimension, a modern imaging protocol with sub-second temporal resolution, and the novel use of multilevel structural equation modeling to demonstrate how neurocomputational signatures of learning can explain associations between stable personality facets and trial-level decision-making. Nonetheless, our observations were cross-sectional in nature and contingent on the naturalistic manipulation of human individual differences, precluding strong causal inferences.

During a social exchange task, we found that learning signals in the default network scaled negatively with callousness, but positively with exploitativeness. Furthermore, default network sensitivity to the accuracy of one’s policy toward a counterpart led to greater social reciprocity in exploitative individuals and lower reciprocity in callous individuals. These results are consistent with evolutionary theories positing that the expansion of the default network coincided with the emergence of sophisticated social cognitive capacities, including counterfactual reasoning and mentalizing. While this development likely contributed to an unprecedented growth in human cooperation, our findings suggest it may have paradoxically contributed to the darker side of human sociality as well, including increased coercion and exploitation.

## Method

For more detail on participants, procedures, and analyses, refer to our Supplemental Materials.

### Participants

The final sample (*n* = 168) included 113 individuals diagnosed with BPD and 55 healthy comparison participants whose behavioral data passed all quality control checks as described in Section I of the Supplemental Materials. Seventeen participants were excluded from imaging analyses due to missing data (*n* = 1), poor quality data (*n* = 4), and excessive motion (*n =* 14; see Supplement Section VII). Participants from the two groups were pooled into a single sample to allow for continuous data analysis across a range of symptom severity. Demographic and clinical characteristics of the final sample are presented in Tables S2-3.

### Experimental Paradigm

Participants completed a modified version of an iterated trust game featuring three different trustees, as depicted in Figure 1B (Vanyukov et al., 2019). Trustees were introduced prior to the task via a picture of a neutral, White, male face from the NimStim Face Stimulus Set (Tottenham et al., 2009) and a short description that included one noteworthy event indicative of the trustee’s “good”, “bad”, or “neutral” reputation^1^. Participants completed a brief questionnaire before and after the task to rate each trustee’s likeability and trustworthiness.

Participants interacted with each trustee for a total of 48 trials. All trustees shared 50% of the time in the first 16 trials, and either 25% (poor) or 88% (rich) of the time in the next 16 trials; the reinforcement schedule reversed in the last 16 trials (i.e., rich-to-poor or poor-to-rich). The order of the trustees was counterbalanced across participants. At the beginning of each trial, participants were told they had $1.00 to either keep or share with the trustee. If they chose to keep, they would retain the $1.00. If they chose to share, they could receive either $1.50 if the trustee also chose to share or be left with nothing if the trustee decided to keep their $1.00.

Earnings from each trustee were shown to participants at the end of each trustee block. Each trial began with a fixation cross, followed by a decision phase, during which participants made their choice to keep or share (decision phase ended after 2.7 seconds when no response was made). The participant’s choice was highlighted for a fixed 200ms, after which participants were shown the trustee’s decision for 1.2 seconds. Critically, participants were shown the trustee’s decision regardless of their own choice, which enabled participants to learn from counterfactual outcomes, with the only difference being that when participants chose to keep, the trustee decision was displayed in gray (as opposed to red or green) to indicate it did not affect earnings for that trial. Intertrial intervals were jittered by combining the unused decision time from the previous decision phase with a base duration sampled from an exponential distribution, with no ITIs allowed to be > 1500ms or < 250ms.

### Factor Analysis

Subscales assessing the agreeableness-antagonism domain were selected from a battery of self-report instruments, including (1) altruism, sympathy, modesty, cooperation, morality, and trust subscales from the *NEO-IPIP-120* (Maples et al., 2014); (2) attention-seeking, callousness, deceitfulness, manipulativeness, and grandiosity subscales from the *Personality Inventory for Diagnostic and Statistical Manual, Fifth Edition* (Krueger et al., 2012); (3) exploitativeness, entitlement rage, and exhibitionism subscales from a modified version of the *Brief Pathological Narcissism Inventory* (Schoenleber et al., 2015); (4) assault, verbal hostility, and negativism subscales from the *Buss Durkee Hostility Inventory* (Buss & Durkee, 1957). Criteria for selecting subscales from each measure are outlined in Supplemental Materials Section V.

Exploratory factor analysis (EFA) using maximum likelihood estimation and an oblimin rotation was conducted on the 17 self-report subscales. One- and two-factor solutions were extracted to attain measures of broadband antagonism and its core facets, respectively. Estimated factor scores from the EFA solutions were saved using the tenBerge method for use in subsequent analyses.

### Behavioral Analysis

Multilevel logistic regression was used to examine the effect of trustee reputation, the trustee’s last decision, and the exchange number with the trustee (i.e., trial within trustee) on participant choices to keep or share. Random slopes were modeled for all three design variables, and a random intercept for subject was included to account for individual differences in cooperation. Age and sex were included as participant-level covariates. These analyses provide an independent corroboration of key behavioral effects relative to the cognitive computational model.

### Computational Modeling

Reinforcement learning models were fit using the variational Bayesian analysis toolbox in MATLAB, which leverages parameter estimates from the full sample to constrain individual parameter estimates, reducing the risk of misestimation in poorly performing participants (Daunizeau et al., 2014). Models employed a version of *Q*-learning with a single hidden state for the expected value of the share action on trial *t*. The value of the keep action was assumed to be updated reciprocally as detailed in our previous study (Vanyukov et al., 2019). The expected value of the share action was updated according to the learning rule:

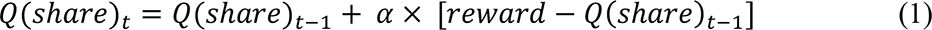

where 𝛼 is the learning rate parameter and 𝑟𝑒𝑤𝑎𝑟𝑑 − 𝑄(𝑠ℎ𝑎𝑟𝑒)_𝑡−1_ is the prediction error.

Choices were modeled using a softmax function that included four free parameters:

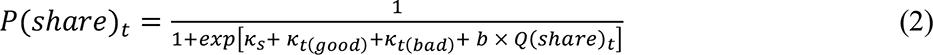

where 𝑏 is a temperature parameter reflecting choice stochasticity and 𝜅_𝑆_ is a participant-level parameter that reflects a general bias to keep or share regardless of reinforcement history. 𝜅_𝑡(𝑔𝑜𝑜𝑑)_ and 𝜅_𝑡(𝑏𝑎𝑑)_ are condition-level parameters that reflect the participant’s bias to keep or share with the good and bad trustees relative to the neutral trustee, respectively. All four competing models included the same free parameters (𝛼, 𝑏, 𝜅_𝑆_, 𝜅_𝑡(𝑔𝑜𝑜𝑑)_, 𝜅_𝑡(𝑏𝑎𝑑)_) but differed with respect to their payout matrices. We have shown in previous work that the models’ parameters are identifiable (Vanyukov et al., 2019). Critically, the distinguishing feature of the policy model was that the valence of the reward on trials in which the participants correctly predicted a trustee defection was positive *relative* to trials in which participants incorrectly predicted a defection.

See Supplemental Section VI for the rationale of competing models and corresponding payout matrices. The winning model was selected by comparing the relative evidence for each model using random effects Bayesian model comparison (BMC), accounting for the full statistical risk incurred (Bayesian omnibus risk [BOR]; Rigoux et al., 2014).

### fMRI Processing and Analysis

Imaging data were acquired in a Siemens Prisma 3T scanner at the Magnetic Resonance Research Center at the University of Pittsburgh. We acquired functional imaging data during the trust task using a simultaneous multislice sequence sensitive to BOLD contrast, TR = .6s, TE = 27ms, flip angle = 45°, multiband acceleration factor = 5, voxel size = 3.1mm^3^. We obtained a sagittal MPRAGE T1 scan, voxel size = 1mm^3^, TR = 2.3s, TE = 3.35ms, GRAPPA 2x acceleration. The anatomical scan was used for coregistration and nonlinear transformation to functional and stereotaxic templates. We also acquired gradient echo fieldmap images (TEs = 4.47 and 6.93ms) for each subject to quantify and mitigate local inhomogeneity of the magnetic field. Neuroimaging data were preprocessed using FMRIPREP version 20.1.1 (Esteban et al., 2019). See Supplement Section VII for full details on preprocessing.

Voxelwise general linear model analyses were conducted using FSL FEAT v6.0 via fmri.factory (Hallquist, 2022a), an R package that streamlines model-based fMRI analyses. Choice and feedback phases of the task were modeled with duration-modulated unit-height boxcar regressors that were convolved with the canonical hemodynamic response function. Participant reaction time on each trial determined the duration of the choice regressor, whereas the duration of the feedback regressor was fixed at 1.10 seconds. Standardized prediction error and value signals derived from the policy model were multiplied by the participant’s decision on each trial (−1/1 = keep/share) to reflect the assumption that policy successes would be associated with greater BOLD activity. Transformed PE and value signals were added as parametric regressors in GLM analyses. PE was aligned to the feedback event and value was aligned to the choice event. To unconfound the influence of response time on BOLD response from the influence of parametric regressors during the choice phase, we convolved the duration-modulated boxcar regressor for the choice event with the HRF, renormalized the peak to 1.0, multiplied the regressor by the corresponding parametric modulator (value), and then summed across trials (cf. Poldrack, 2015). Trustee type was included as a within-subjects factor, allowing for linear contrasts among trustees. Non-response trials were modeled independently from trials on which participants made responses.

Group-level analyses were carried out using FSL FLAME 1+2 with automatic outlier deweighting (Woolrich, 2008), which implements Bayesian mixed effects estimation of the group parameter estimates including full Markov Chain Monte Carlo-based estimation for near-threshold voxels (Woolrich et al., 2004). Group analyses included age and sex as covariates of no interest. To correct for familywise error at the whole-brain level, we applied the probabilistic threshold-free cluster enhancement method (pTFCE), thresholding whole-brain maps at *FWE p* < .05 (Spisák et al., 2019). This algorithm provides strict control over familywise error and boosts sensitivity to clusters of activated voxels.

To examine how individual differences in default network BOLD response related to learning and behavior, we extracted participant-level regression coefficients (“betas”) from the prediction error contrast of the whole-brain fMRI GLM analyses based on an a priori specified atlas that combined leading cortical and subcortical parcellations (Choi et al., 2012; Hall & Hallquist, 2022; Schaefer et al., 2018). The atlas included 244 (200 cortical + 44 subcortical) parcels assigned to 17 functional networks (Yeo et al., 2011). In total, 37 parcels were assigned to one of the three default network subsystems in Yeo et al. (2011). All voxels within each DN parcel were included in the extracted betas. To avoid redundancy, fMRI betas were extracted from an intercept-only group model, as age and sex were entered in covariates in all MSEM models. To derive an index of PE-modulation of the default network, we entered betas from all parcels assigned to the default network into a factor analysis, extracted a one-factor solution, and saved estimated factor scores using the tenBerge method (Ten Berge et al., 1999). Likewise, to derive indices of PE modulation of each default network subsystem, we entered betas only from parcels assigned to each subsystem into a factor analysis (three total), extracted a one-factor solution from each, and saved estimated factor scores. The advantage of using factor analyses to derive these indices (as opposed to using a mean composite) is that it allows for individual parcels to be weighted based on their respective contribution to the broader network. Factor scores derived with this method were comparable to those using a more data-driven approach in which all parcels were included and additional factors were extracted to derive subsystem indices (i.e., we extracted 1, 2, 3, and 4-factor solutions using all parcels; see Supplemental Section IX).

### Multilevel Structural Equation Modeling

Relations between personality, learning signals encoded in the brain, and trial-level behavioral performance were tested using multilevel structural equation modeling (MSEM), which enables the simultaneous examination of multiple predictors and outcomes in hierarchically structured data (Sadikaj et al., 2019). In MSEM, latent decomposition is used to partition total variance in outcomes and predictors into within- and between-person components All MSEM models were estimated using Bayesian estimation with noninformative priors in *Mplus Version 8.7* (Muthén & Muthén, 2019). Bayesian estimation uses all available data and provides similar results to full information maximum likelihood in accounting for missing data (Asparouhov & Muthén, 2010). For all MSEM models, we report unstandardized and standardized regression coefficients, 95% credible intervals, and Bayesian *p* values. Bayesian *p* values are based on the probability of direction test, a hypothesis test that is closely aligned to frequentist null hypothesis significance testing (Makowski et al., 2019). Note, however, that Bayesian posterior probabilities quantify the extent to which the data support a given hypothesis, which provides stronger inference than frequentist approaches that quantify the probability of observing the data under the null hypothesis.

In all MSEM models, random slopes were specified for the effects of trustee reputation, trustee’s previous decision, and trial within trustee on participant decisions to keep or share (Bolger et al., 2019). To examine the effect of learning signals on trial-level behavior, default network PE factor scores were added as between-subjects moderators of design effects (see Figure 1D). The ability of MSEM to accommodate multiple outcomes allows for formal tests of mediation across hierarchical levels in the data. In models examining associations between personality, learning signals, and behavior, personality traits were added as between-subjects predictors of default network learning signals. Mediation (the effect of personality on trial-level behavior via learning signals) was tested using a product of coefficients approach. In contrast to conventional mediation analysis, Bayesian mediation analysis constructs credible intervals for indirect effects that are not subject to normality assumptions on the sampling distribution of the estimate of the indirect effect, and which do not rely on large sample approximations, allowing for more exact inferences (Yuan & MacKinnon, 2009). All models controlled for age and sex as between-subjects covariates.

## Acknowledgments

This work was supported by the National Institute of Mental Health [K01-MH123915 (TAA); R01-MH048463 (AYD & MNH); R01-MH119399 (MNH)]. The authors would like to thank Jacob Koudys, Vanessa Brown, and Polina Vanyukov for assistance preparing the manuscript, refining code, and offering feedback on earlier versions of this manuscript.

## Author Contributions

TAA conducted all analyses, developed primary hypotheses, and wrote the manuscript. AYD and MNH designed the experimental task and computational model, developed the codebase, oversaw data collection, and provided revisions to the manuscript.

## Footnotes

1 58 participants also interacted with a “computer” trustee. For the purposes of this analysis, trials involving the computer trustee were treated as missing.

