## Supplemental Appendix for "The Dark Side of Mentalizing: Learning Signals in the Default Network During Social Exchanges Support Cooperation and Exploitation"

**Table of Contents**

**Section I:** Sampling and Recruitment, Missing Data, and Quality Control Exclusions
**Section II**: Clinical and Demographic Characterization
**Section III:** Manipulation Checks
**Section IV:** Behavioral Results
**Section V:** Measure Selection & Exploratory Factor Analysis
**Section VI:** Computational Modeling
**Section VII**: Neuroimaging Preprocessing
**Section VIII**: Whole Brain Prediction Error Clusters
**Section IX**: Comparison of *a priori* derived Default Network Subsystems to Data-Driven Default Network Factors
**Section X:** Effect of Prediction Error Signaling in the Default Network on Behavior
**Section XI.** Association between Personality and Prediction Error Modulation in the Default Network.
**Section XII.** Effect of Callousness and Exploitativeness on Behavior via Prediction Error Modulation of the Default Network
**Section XIII.** Direct Effects of Personality on Behavior
**Section XIV.** Effect of Antagonism on Behavior via Prediction Errors in the Default Network

**Section I. Sampling and Recruitment, Missing Data, and Quality Control Exclusions**

Participants were recruited via community advertisements and referrals from local inpatient and outpatient clinics. Exclusionary criteria included a lifetime diagnosis of psychosis or bipolar disorder, clinical evidence of organic brain disease, physical ailments with known psychiatric consequences, and IQ < 70 as measured by the Wechsler Test of Adult Reading (Wechsler, 2001). Participants diagnosed with BPD had to meet probable or definite criteria for a lifetime diagnosis based on the International Personality Disorder Examination [IPDE; (Loranger et al., 1987)]. Healthy comparison participants were required to have no lifetime Axis I or II diagnoses as determined by the Structured Clinical Interview for DSM-IV or IPDE.

In total, 177 participants completed the iterative social trust game. Of these, seven subjects were excluded for pressing the same button throughout the entire task. One subject was excluded for only responding on 12% of trials. One additional subject was excluded due to a technical difficulty that resulted in the participant viewing incorrect trustee stimuli. This yielded a final behavioral sample of 168 individuals. Due to a technical glitch, the task ended one trial early for three participants. Responses on these trials were treated as missing in behavioral analyses, and the trials were dropped from imaging analyses.

Of the 168 participants with behavioral data, one participant did not have imaging data, and three others were excluded from the imaging analyses due to poor quality data. Fourteen additional subjects were excluded due to head motion (details in Section VII below), resulting in a final fMRI sample size of 150.

Of the 168 participants with behavioral data, 161 had at least one self-report measure, as can be seen in the table below.

Table S1. *Rates of Missing Data*

| Measure | Sample Size |
|---|---|
| Modified Social Trust Game | 168 |
| Pre/Post Trustee Ratings | 149 |
| <u>Self-Reports</u> | 161 |
| Buss-Durkee Hostility Inventory | 157 |
| Personality Inventory for DSM-5 | 158 |
| NEO-IPIP-120 | 158 |
| Brief Pathological Narcissism Inventory | 153 |
| <u>fMRI</u> | 150 |

**Section II. Overview of the Sample**

Table S2. *Demographic Characterization of the Sample*

| Demographic Variable | |
|---|---|
| Mean Age (SD) | 31.16 (9.30) |
| Female (%) | 131 (78%) |
| Mean Years of Education (SD) | 15.46 (2.20) |
| N Race (%) | |
| American Indian | 4 (2.38%) |
| Asian | 16 (9.52%) |
| Black | 26 (15.48%) |
| Native Hawaiian/Pacific Islander | 3 (1.79%) |
| White | 134 (79.76%) |
| Prefer not to answer | 5 (2.98%) |
| N Marital Status (%) | |
| Never Married | 108 (66.26%) |
| Married | 27 (16.56%) |
| Separated | 3 (1.84%) |
| Divorced | 14 (8.59%) |
| Widowed | 1 (.61%) |
| Cohabitating | 8 (4.91%) |
| Pending Marriage | 2 (1.23%) |
| Previous Suicide Attempt | 84 (50.00%) |

Table S3. *Clinical Characterization of the Sample at Study Enrollment*

| Diagnosis (Lifetime) | N (%) |
|---|---|
| Agoraphobia | 5 (3.31%) |
| Anorexia | 8 (5.30%) |
| Binge Eating | 13 (8.61%) |
| Body Dysmorphia (Current Only) | 4 (2.65%) |
| Bulimia Nervosa | 15 (9.93%) |
| Dysthymia | 5 (3.31%) |
| Generalized Anxiety (Current Only) | 63 (41.72%) |
| Hypochondriasis | 1 (.66%) |
| Major Depression | 91 (60.26%) |
| Obsessive-Compulsive | 18 (11.92%) |
| Panic | 42 (27.81%) |
| Post-traumatic Stress | 54 (35.76%) |
| Social Phobia | 37 (24.5%) |
| Specific Phobia | 31 (20.53%) |
| Other DSM Axis-I | 3 (1.99%) |
| <u>Substance Use Disorders</u> | |
| Alcohol | 55 (36.42%) |
| Cannabis | 30 (19.87%) |
| Cocaine | 12 (7.95%) |
| Hallucinogens/PCP | 3 (1.99%) |
| Opioid | 9 (5.96%) |
| Polysubstance | 10 (6.62%) |
| Sedatives/Hypnotics/Anxiolytics | 7 (4.64%) |
| Stimulants | 6 (3.97%) |
| <u>Personality Disorders</u> | |
| Antisocial | 14 (8.33%) |
| Avoidant | 26 (15.48%) |
| Borderline | 113 (67.26%) |
| Dependent | 39 (23.21%) |
| Histrionic | 9 (5.36%) |
| Narcissistic | 1 (.60%) |
| Obsessive-Compulsive | 15 (8.93%) |
| Paranoid | 18 (10.71%) |
| Schizoid | 1 (.60%) |
| Schizotypal | 3 (1.79%) |

*Note.* IPDE $n$ = 168. SCID $n$ = 151. SCID / IPDE interviews were completed at enrollment of a longitudinal study. Task data was collected at enrollment for new participants or at a scheduled reassessment for participants already enrolled in the longitudinal component of the study.

**Section III. Manipulation Checks: Trustworthiness and Likeability Ratings**

To examine the effect of our reputation manipulation, trustee (good, bad, neutral) ratings on trustworthiness and likeability pre- and post-task were entered into a 3 × 2 repeated measures ANOVA ($n$ = 149 due to missingness on survey data). Ratings differed significantly across trustees (*trustworthiness:* $F_{(1.71, 253.31)} = 149.37$, $p < .001$; *likeability:* $F_{(1.75, 259.56)} = 80.42$, $p < .001$) and time (*trustworthiness:* $F_{(1, 148)} = 19.12$, $p < .001$; *likeability:* $F_{(1, 148)} = 35.30$, $p < .001$). Post hoc comparisons using Bonferroni correction indicated that on average, participants rated bad trustees as less trustworthy and likeable than good (*trustworthiness:* $t = -14.97$, $p < .001$; *likeability:* $t = -12.59$, $p < .001$) and neutral (*trustworthiness:* $t = -10.25$, $p < .001$; *likeability*: $t = -7.04$, $p < .001$) trustees, and neutral trustees as less trustworthy and likeable than good (*trustworthiness:* $t = -7.96$, $p < .001$; *likeability:* $t = -7.70$, $p < .001$) trustees. Both sets of ratings also declined from pre- to post-task, on average (*trustworthiness:* $t = -3.77$, $p < .001$; *likeability:* $t = -5.34$, $p < .001$).

Two-way time × trustee interactions (*trustworthiness*: $F_{(2, 296)} = 38.61$, $p < .001$; *likeability*: $F_{(2, 296)} = 18.89$, $p < .001$) indicated that differences between pre- and post-task ratings varied by trustee. Specifically, ratings of trustworthiness and likeability decreased pre- to post-task for the good and neutral trustees. In contrast, trustworthiness ratings increased from pre- to post-task for the bad trustee (there was no difference in likeability ratings for the bad trustee before and after the task following Bonferroni correction).

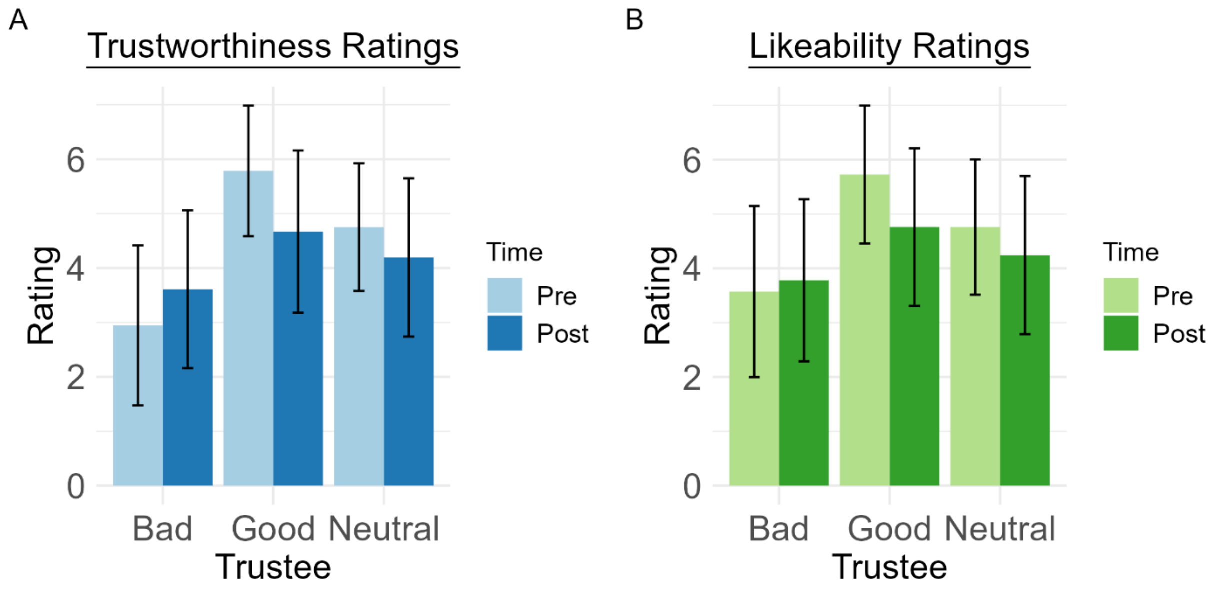

**Figure S1.** Pre- and post-task ratings of trustee trustworthiness and likeability.

**Section IV. Behavioral Results**

To examine behavioral performance on the trust game, we fit a random intercept-random slope model with fixed effects of reputation, the trustee's decision on the preceding trial, and the exchange number (i.e., within-trustee trial number). Random slopes were modeled for all three design variables.

Table S4. *Behavioral Effects on Modified Social Trust Game.*

| | *β (B)* | *SE* | *p* |
|---|---|---|---|
| Fixed Effects | | | |
| Intercept | --- (-.15) | .15 | .32 |
| Age | .70 (.35) | .14 | < .001 |
| Sex (0 = M, 1 = F) | -.15 (-.18) | .14 | .27 |
| Exchange Number | -.14 (-.05) | .03 | < .001 |
| Bad Reputation | -.24 (-.26) | .05 | < .001 |
| Good Reputation | .02 (.02) | .05 | .64 |
| Trustee Decision (t-1) | .62 (.62) | .07 | < .001 |
| Random Effects | *Variance* | *SE* | |
| Intercept (subject) | .94 | .97 | |
| Exchange Number | .00 | .07 | |
| Bad Reputation | .31 | .56 | |
| Good Reputation | .23 | .48 | |
| Trustee Decision (t-1) | .78 | .88 | |

*Note*. Between-subjects $n = 168$; within-subjects $n = 23369$.

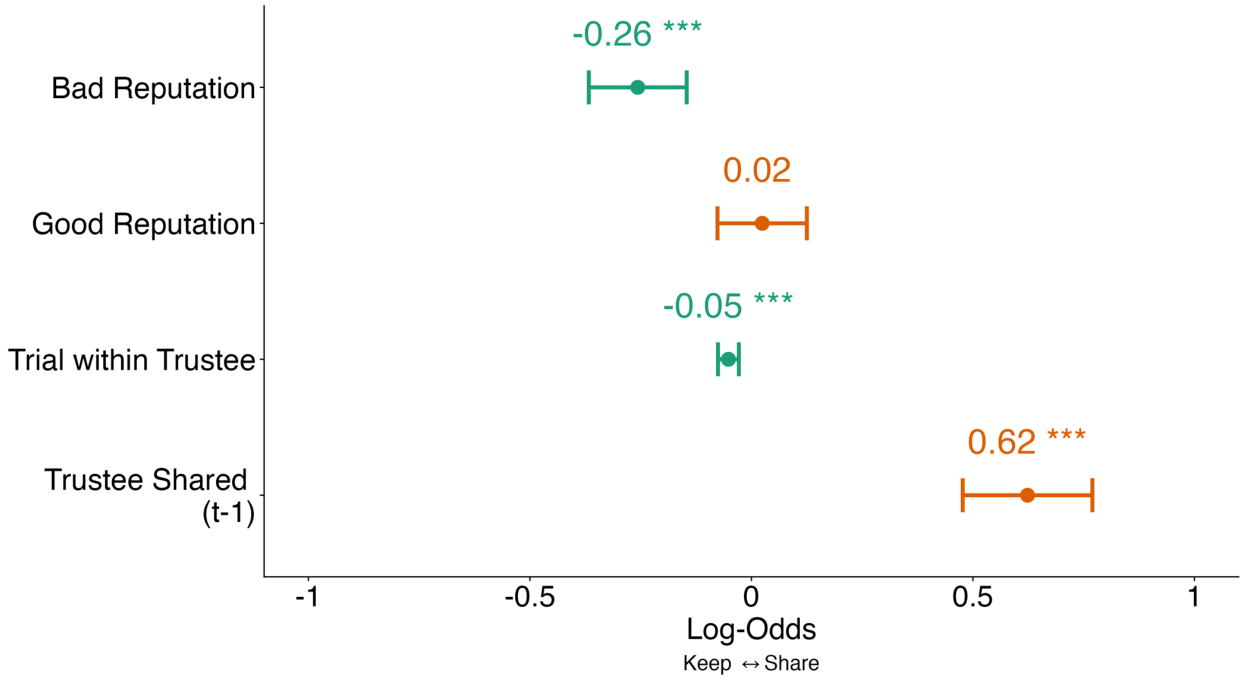

**Figure S2.** Effect of task manipulations on participants' decision to keep (coded 0) or share (coded 1).

**Section V. Measure Selection & Exploratory Factor Analysis**

We conducted exploratory factor analysis (EFA) using 17 self-report scales selected a priori as central to the antagonism domain (see below for criteria used to select scales). A one-factor solution indicated that most scales exhibited moderate to strong loadings on a general antagonism factor (*M* loading = .63, range = [-.09, .89]). Antagonism accounted for 45% of the variance among all indicators. Velicer's MAP(Velicer, 1976) test indicated that a two-factor solution to the EFA was optimal, consistent with structural research showing the antagonism-agreeableness domain can be divided into two correlated lower-order facets (Allen et al., 2019). In the two-factor solution, the first factor, labeled exploitativeness, was marked by scales measuring grandiosity, dishonesty, deceitfulness, and manipulativeness. The second factor, labeled callousness, was defined by scales related to aggression and lack of empathy. The two factors were correlated at $r = .53$ and accounted for 29% and 26% of variance among indicators, respectively. Factor scores from the one- and two-factor solutions were saved using the tenBerge method for use in subsequent analyses.

Table S5. *Exploratory Factor Analysis of Antagonism-Agreeableness Scales*

| | One Factor Solution | Two-Factor Solution | |
|---|---|---|---|
| Subscale | Antagonism | Exploitativeness | Callousness |
| Attention-Seeking | .75 | **.97** | -.15 |
| Exhibitionism | .78 | **.95** | -.09 |
| Manipulativeness | .89 | **.69** | .27 |
| Morality | -.86 | **-.65** | -.28 |
| Deceitfulness | .87 | **.63** | .34 |
| Exploitativeness | .83 | **.62** | .28 |
| Entitlement Rage | .79 | **.59** | .31 |
| Modesty | -.39 | **-.55** | .14 |
| Grandiosity | .67 | **.40** | .39 |
| Callousness | .72 | .11 | **.81** |
| Cooperation | -.74 | -.18 | **-.74** |
| Trust | -.45 | .09 | **-.69** |
| Altruism | -.32 | .23 | **-.68** |
| Assault | .54 | .06 | **.65** |
| Verbal Aggression | .50 | .05 | **.61** |
| Negativism | .51 | .19 | **.42** |
| Sympathy | -.09 | .22 | **-.37** |

*Note.* $n = 161$. Higher indicator loading between the two facets are bolded.

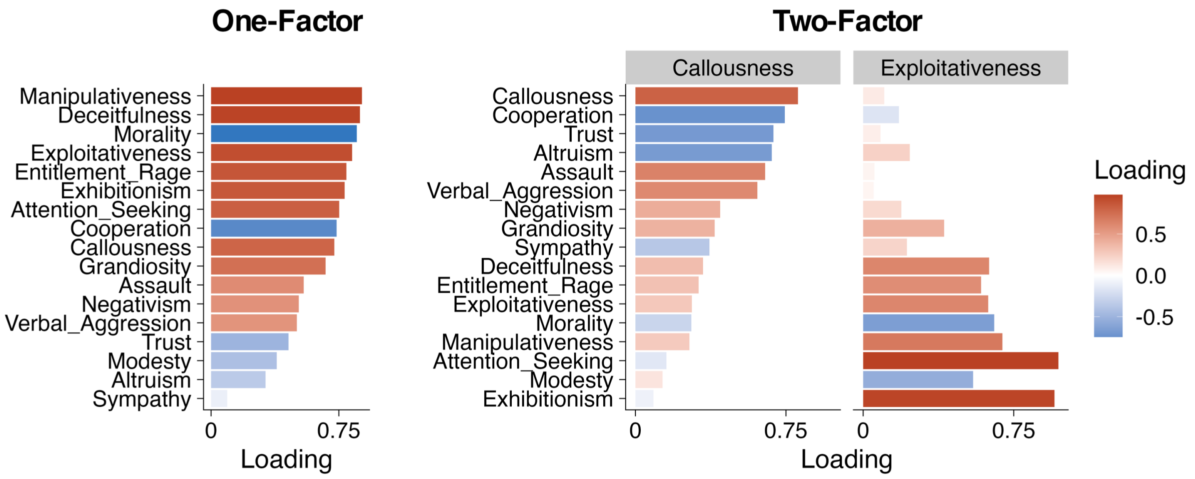

**Figure S3.** Exploratory factor analysis of 17 self-report scales related to the Antagonism v. Agreeableness domain.

*Subscale Selection Criteria*

Subscales from each of the self-report measures were selected for inclusion in the EFA based on the following criteria:

1. For the NEO-IPIP-120, all facets assigned to the Agreeableness domain were included (Trust, Morality, Modesty, Sympathy, Altruism, Cooperation)
2. For the PID-5, all facets with a primary loading on Antagonism in Krueger et al., 2012 were included (Manipulativeness, Attention-Seeking, Deceitfulness, Grandiosity, Callousness).
3. For the Buss-Durkee Hostility Inventory and the Brief Pathological Narcissism Inventory, a subscale was included if it a) was more strongly correlated with IPIP-Agreeableness than any of the other Big Five domains on the IPIP; and b) did not exhibit a cross-correlation with any other IPIP domain within .10 of its correlation with Agreeableness (Buss-Durkee: Assault, Verbal, Negativism; BPNI: Exploitativeness, Entitlement Rage, Exhibitionism). This yielded 17 total self-report scales across the four measures.

*Note on Modified BPNI*

We used a modified version of the BPNI scale that adds four items to comprise an exhibitionism scale. The items were:

I like to be the center of attention.
I will usually show off if I get the chance.
I do things to draw attention to myself on purpose.
I sometimes do things just so that people will be sure to notice me.

**Section VI. Computational Modeling**

We have shown in previous work that the policy model dominates alternative accounts of behavior on the modified social trust game (Vanyukov et al., 2019). Simulations performed as part of our earlier study indicated that alternative models tested herein were distinguishable from one another (only the model that generated a dataset provided the best fit to that dataset) and all model parameters were identifiable. The current paper builds on these basic validations of the model and focuses on how learning signals derived from the policy model relate to social exchanges and individual differences in personality. As such, we encourage readers to review prior parameter recovery and model identifiability analyses in Vanyokov et al., 2019.

In the current analyses, we tested four alternative accounts of how participants represent reinforcement on the modified trust game, including: 1) an actual rewards model, in which participants learned only from experienced outcomes on the task; 2) a regret model, in which participants tracked the difference between the experienced and best possible outcome; 3) a counterpart-oriented counterfactual model, in which participants track the difference between the experienced outcome and what the outcome would have been had the counterpart acted differently; and 4) the policy model, in which participants track the difference between the experienced outcome and, for their untaken actions, what the outcome would have been had they acted differently. As outlined in the main text, all four alternative accounts included the same four model parameters, but varied in how reinforcement was encoded according to the following payout matrices:

*Payout Matrices for Alternative Models*

| Model | Subject Invests / Trustee Returns | Subject Invests / Trustee Keeps | Subject Keeps / Trustee Returns | Subject Keeps / Trustee Keeps |
|---|---|---|---|---|
| Actual Rewards | 1.5 | 0 | 1 | 1 |
| Regret | 1.5 | -1.5 | -.5 | -.5 |
| Counterpart Counterfactual | 1.5 | -1.5 | 0 | 0 |
| Policy | 1.5 | 0 | -.5 | 1 |

All models were fit using Variational Bayesian Analysis in MATLAB (Daunizeau et al., 2014). The variational Bayesian approach leverages parameter estimates from the entire sample to constrain estimates for individuals, reducing the risk of misestimation in poorly performing subjects. This yields more robust and precise estimates of individual parameters and evolving hidden states.

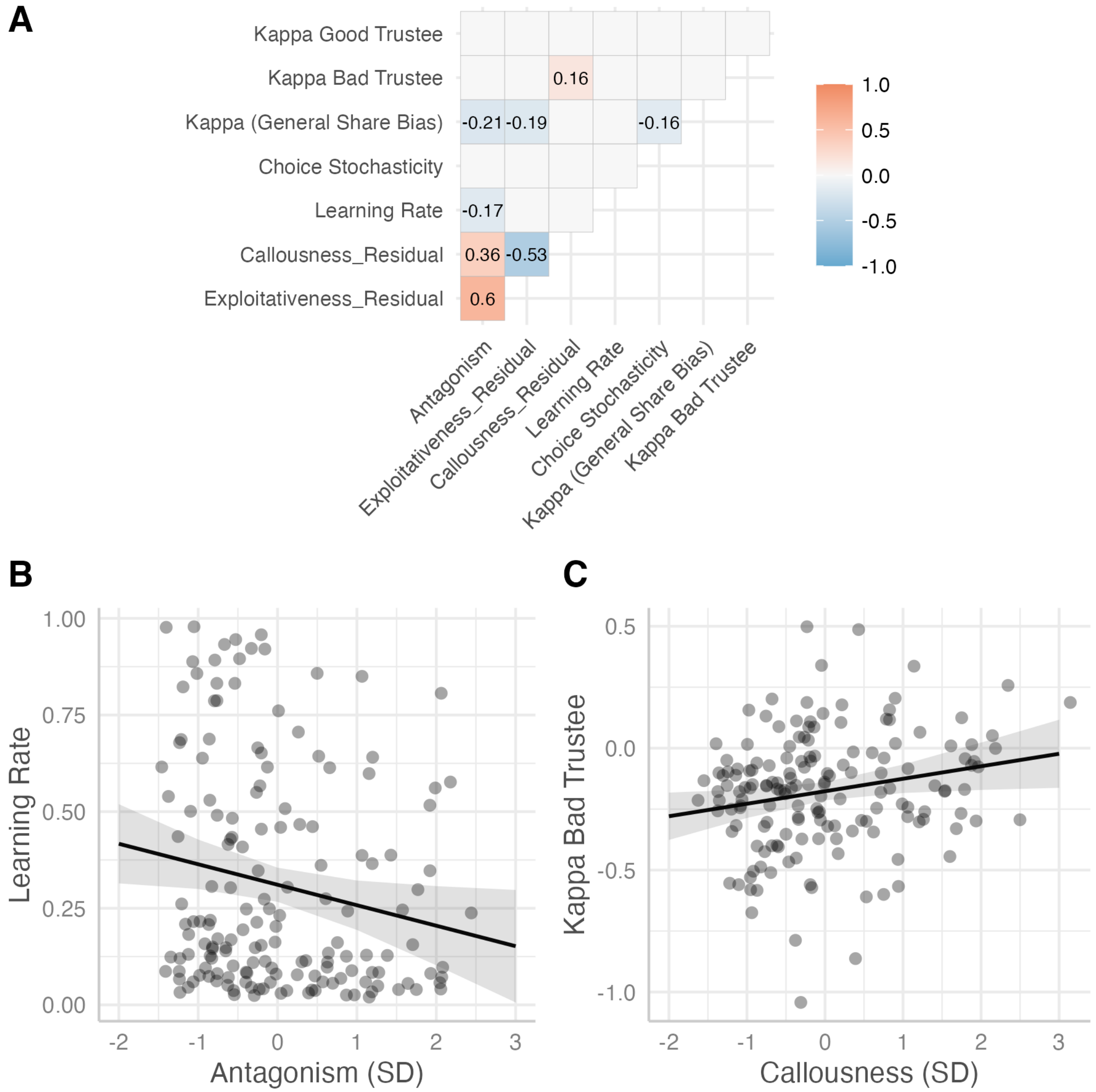

**Figure S4.** A) Correlation plot of the association between traits and model parameters (only displaying correlations significant at $p < .05$). Callousness and exploitativeness represent residuals after controlling for shared variance with the other facet (i.e., each facet was regressed on the other, and the residuals were saved). This has the effect of partialing out variance shared between the facets, or variance that contributes to antagonism. Variances unique to callousness and exploitativeness, and independent of antagonism, are negatively correlated with each other as seen in the corrplot above. B) Antagonism negatively predicted the learning rate after controlling for age and sex. C) Callousness positively predicts the $\kappa$ parameter for the bad trustee after controlling for age, sex, and exploitativeness.

**Section VII. Neuroimaging Preprocessing**

MRI preprocessing was performed using FMRIPREP version 20.1.1 (Esteban et al., 2019), a Nipype (Gorgolewski et al., 2011) based tool. Each T1-weighted volume was corrected for INU (intensity non-uniformity) using N4BiasFieldCorrection v2.1.0 (Tustison et al., 2010) and skull-stripped using antsBrainExtraction.sh v2.1.0 (using the OASIS template). Brain surfaces were reconstructed using recon-all from FreeSurfer v6.0.1 (Dale et al., 1999), and the brain mask estimated previously was refined with a custom variation of the method to reconcile ANTs-derived and FreeSurfer-derived segmentations of the cortical gray-matter of Mindboggle (Klein et al., 2017). Spatial normalization to the ICBM 152 Nonlinear Asymmetrical template version 2009c (Fonov et al., 2009) was performed through nonlinear registration with the *antsRegistration* tool of *ANTs* v2.1.0 (Avants et al., 2008), using brain-extracted versions of both T1w volume and template. Brain tissue segmentation of cerebrospinal fluid (CSF), white-matter (WM) and gray-matter (GM) was performed on the brain-extracted T1w using fast FSL v5.0.9 (Zhang et al., 2001).

Functional data was slice time corrected using *3dTshift* from AFNI v16.2.07 (Cox, 1996) and motion corrected using *mcflirt* (FSL v5.0.9; Jenkinson et al., 2002). Distortion correction was performed using fieldmaps processed with fugue (Jenkinson, 2003). This was followed by co-registration to the corresponding T1w using boundary-based registration (Greve & Fischl, 2009) with six degrees of freedom using bbregister (FreeSurfer v6.0.1). Motion correcting transformations, BOLD-to-T1w transformation and T1w-to-template (MNI) warp were concatenated and applied in a single step using *antsApplyTransforms* (ANTs v2.1.0) using Lanczos interpolation. Images were spatially smoothed using a 6mm full-width at half maximum (FWHM) kernel.

Framewise displacement (Power et al., 2014) was calculated for each functional run using the implementation of Nipype. To reduce head motion artifacts, we then conducted an independent component analysis for each run using FSL MELODIC. The spatiotemporal components were then passed to a classification algorithm, ICA-AROMA, validated to identify and remove motion-related artifacts (Pruim et al., 2015). Components identified as noise were regressed out of the data using FSL regfilt (non-aggressive regression approach). ICA-AROMA has performed very well in head-to-head comparisons of alternative strategies for reducing head motion artifacts (Ciric et al., 2017). We then applied a .008 Hz temporal high-pass filter to remove slow-frequency signal changes (Woolrich et al., 2001); the same filter was applied to all regressors in GLM analyses. Finally, we renormalized each voxel time series to have a mean of 100 to provide similar scaling of voxelwise regression coefficients across runs and participants.

In addition to mitigating head motion-related artifacts using ICA-AROMA, we excluded subjects who had runs in which more than 15% of volumes had a framewise displacement of .5 mm or greater, as well as runs that had a maximum framewise displacement greater than 5mm at any point in acquisition. Additionally, in voxelwise GLMs, we included the mean time series from the cerebral white matter and ventricles, as well as the derivatives of these signals, as confound regressors (Ciric et al., 2017).

**Section VIII. Whole Brain Prediction Error Clusters**

Table S6. *PTFCE-Corrected Prediction Error Map*

| ROI Number | Number of Voxels | Maximum Intensity | Center-of-Mass MNI Coordinates | | | Regions |
|---|---|---|---|---|---|---|
| | | *z*-max | x | y | z | |
| 1 | 19717 | 15.318 | -15.2 | 8.1 | -10.3 | |
| 1.1 | 1445 | 15.318 | -15.2 | 8.1 | -10.3 | Bilateral Putamen, Bilateral Striatum, Bilateral Hippocampus, Bilateral Amygdala |
| 1.2 | 1093 | 12.264 | -55.9 | -66.9 | -10.3 | Left Lateral Occipital Cortex, Left Inferior Temporal Gyrus, Left Middle Temporal Gyrus |
| 1.3 | 1056 | 13.593 | 6.6 | -45 | 33.1 | Precuneus, Posterior Cingulate, Anterior Cingulate |
| 1.4 | 951 | 11.51 | 50.4 | -60.6 | -13.4 | Right Lateral Occipital Cortex, Right Inferior Temporal Gyrus, Right Middle Temporal Gyrus |
| 1.5 | 882 | 12.638 | -49.6 | -38.8 | 61 | Left Postcentral Gyrus, Left Supramarginal Gyrus, Left Precentral Gyrus |
| 1.6 | 616 | 12.108 | 37.9 | -70 | -41.3 | Right Cerebellum |
| 1.7 | 455 | 12.242 | -46.5 | -73.1 | -35.1 | Left Cerebellum |
| 1.8 | 394 | 12.441 | 0.4 | 51.9 | -10.3 | Frontal Pole, Frontal Medial Cortex, Paracingulate Gyrus |
| 2 | 594 | -11.347 | 9.8 | 14.4 | 70.3 | Superior Frontal Gyrus, Supplementary Motor Area, Paracingulate Gyrus, Anterior Cingulate Gyrus |
| 3 | 164 | 7.3205 | 25.4 | 33.1 | 51.7 | Right Superior Frontal Gyrus |
| 4 | 70 | -6.067 | 19.1 | -60.6 | 8.3 | Right Precuneus, Right Lingual Gyrus, Right Intracalcerine Cortex |
| 5 | 70 | -6.5642 | 31.6 | 20.6 | 8.3 | Right Insular Cortex, Right Frontal Operculum |
| 6 | 46 | -6.2434 | -43.4 | 14.4 | 5.2 | Left Frontal Operculum |
| 7 | 30 | -5.7791 | 47.2 | -13.8 | 48.6 | Right Postcentral Gyrus, Right Precentral Gyrus |
| 8 | 20 | -6.2118 | 50.4 | 8.1 | -38.2 | Right Temporal Pole |
| 9 | 18 | -5.7903 | -40.2 | 20.6 | -13.4 | Left Frontal Orbital Cortex |
| 10 | 17 | -6.0904 | -21.5 | -51.2 | 2.1 | Left Lingual Gyrus |

*Note*. Threshold $z_{ptfce} > 5.16$; FWER $p_{ptfce} < .05$. Clusters > 2000 voxels were broken up into subclusters of at least 100 voxels to ease interpretation and region labeling. Age and sex were included as group-level covariates.

**Section IX. Comparison of *a priori* derived Default Network Subsystems to Data-Driven Default Network Factors**

Meta-analytic research on the default network has described it as a functionally heterogenous network that spans three subsystems: the core, dorsal medial, and medial temporal subsystems (Andrews-Hanna et al., 2014). In our primary analysis, we ensured consistency with the extant literature on the default network by using an atlas to extract regression coefficients ("betas") from parcels assigned to each subsystem based on the parcellation of Schaefer and colleagues (2018). Subsequently, we conducted a series of factor analyses to derive factor scores that were used to represent PE modulation within each subsystem. One drawback to this approach is that PE signaling across parcels of the default network in our study may not exhibit a covariance structure that adheres to the canonical medial temporal, core, and dorsal medial subsystems. To address this possibility, we took a data-driven approach to deriving default network subsystems by including all 37 default network parcels in a factor analysis employing maximum likelihood estimation and an oblimin rotation. The three-factor solution yielded factors resembling the medial temporal subsystem, the anterior hub of the default network, and a final factor that blended the dorsal medial subsystem and posterior hub. We also examined a four-factor solution to determine whether the posterior hub would diverge from the dorsal medial subsystem. Indeed, the four-factor solution yielded clear factors representing the medial temporal and dorsal medial subsystems, along with the anterior and posterior cores. As seen below, these data-derived factors were strongly correlated with the a priori derived subsystems used in our main analysis, though the a priori derived core subsystem was weighted toward the posterior hub as opposed to the anterior hub. In general, there was a high degree of correlation across data-driven and a priori factors, suggesting the method used to derive subsystem factors did not qualify our results.

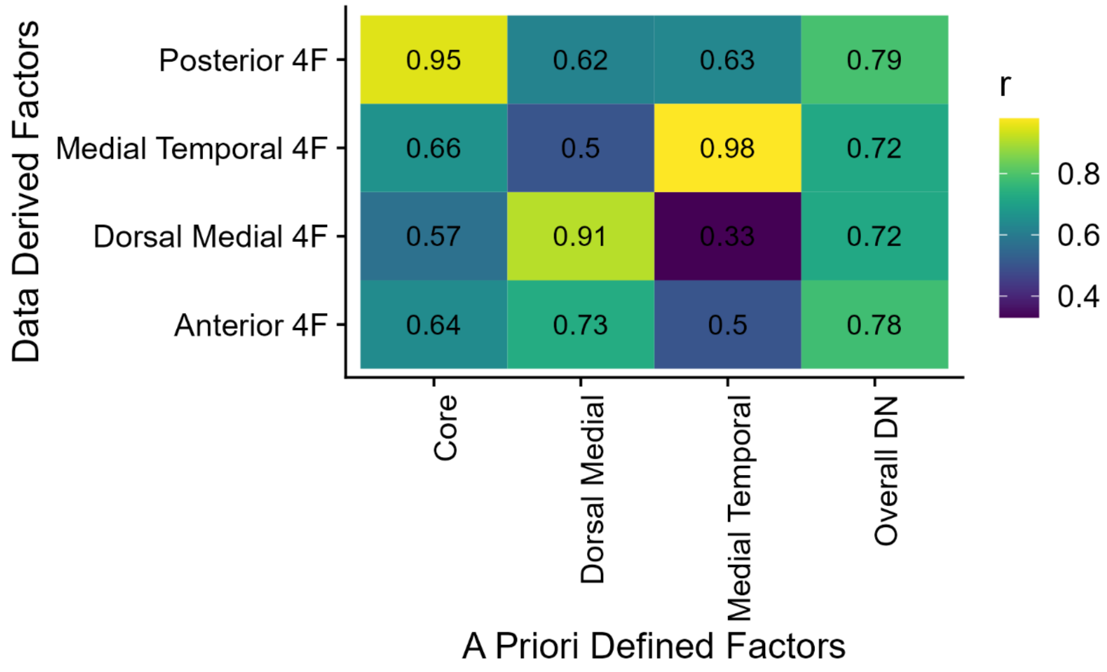

**Figure S5.** Correlations between data-derived and a priori defined factors.

**Section X. Effect of Prediction Error Signaling in the Default Network on Behavior**

Table S7. *Multilevel Structural Equation Model Examining Moderation of the Effects of Reputation and Reciprocity on Decision to Share by Prediction Errors in the Default Network*

| Variable | *β (B)* | *95% CI* | *p* |
|---|---|---|---|
| Between-Person | | | |
| Share Intercept | --- (.04) | -.12, .23 | .64 |
| Share on PE-DN | .12 (.07) | (-.06, .27) | .17 |
| Share on Age | .23 (.18) | (.12, .32) | < .001 |
| Share on Sex | -.04 (-.08) | (-.15, .06) | .44 |
| $a_i$ on PE-DN | -.01 (-.00) | (-.17, .19) | .96 |
| $b_i$ on PE-DN | -.12 (-.04) | (-.33, .11) | .30 |
| $c_i$ on PE-DN | .25 (.14) | (.09, .39) | < .001 |
| $a_i$ on Age | -.04 (-.01) | (-.16, .08) | .54 |
| $b_i$ on Age | -.11 (-.05) | (-.25, .02) | .09 |
| $c_i$ on Age | -.06 (-.05) | (-.18, .05) | .31 |
| $a_i$ on Sex | -.03 (-.01) | (-.15, .10) | .66 |
| $b_i$ on Sex | -.03 (-.03) | (-.16, .12) | .69 |
| $c_i$ on Sex | .05 (.09) | (-.07, .15) | .44 |
| Within-Person | | | |
| Share on Exchange ($a_i$) | -.05 (-.03) | (-.06, -.03) | < .001 |
| Share on Reputation ($b_i$) | -.08 (-.16) | (-.09, -.06) | < .001 |
| Share on Trustee Decision (*t-1*) ($c_i$) | .16 (.31) | (.14, .18) | < .001 |

*Note.* Except for the intercept of the decision to share, all standard deviations and credibility intervals refer to standardized regression coefficients. Italicized *a, b,* and *c* parameters represent random slopes included in the model. In MSEM, random slopes reflect between-person differences in the strength of a within-person effect (e.g., $b_i$ captures individual differences in the effect of reputation on decisions to share), which can then be related to other between-person predictors. Correspondence between random slopes and within-person associations is denoted in the parentheticals in the Within-Person section of the table. Within-person $n$ = 24192. Between-persons $n$ = 168.

Table S8. *Multilevel Structural Equation Model Examining Moderation of Reputation and Reciprocity Effects on Decision to Share by Prediction Errors in Default Network Subsystems*

| Variable | *β (B)* | *95% CI* | *p* |
|---|---|---|---|
| <u>Between-Person</u> | | | |
| Share Intercept | --- (.05) | -.12, .25 | .48 |
| Share on PE-Core | .06 (.04) | (-.21, .32) | .69 |
| Share on PE-Medial Temporal | .10 (.06) | (-.11, .33) | .42 |
| Share on PE-Dorsal Medial | -.03 (-.02) | (-.24, .20) | .82 |
| Share on Age | .23 (.19) | (.13, .32) | < .001 |
| Share on Sex | -.05 (-.09) | (-.15, .06) | .40 |
| $a_i$ on PE-Core | .00 (-.00) | (-.26, .29) | .99 |
| $b_i$ on PE-Core | -.03 (-.01) | (-.35, .29) | .93 |
| $c_i$ on PE-Core | .05 (.03) | (-.24, .33) | .67 |
| $a_i$ on PE-Medial Temporal | .15 (.02) | (-.11, .38) | .30 |
| $b_i$ on PE-Medial Temporal | -.31 (-.11) | (-.53, -.03) | .04 |
| $c_i$ on PE-Medial Temporal | .17 (.09) | (-.10, .37) | .13 |
| $a_i$ on PE-Dorsal Medial | -.09 (-.01) | (-.33, .14) | .47 |
| $b_i$ on PE-Dorsal Medial | .18 (.06) | (-.13, .39) | .24 |
| $c_i$ on PE-Dorsal Medial | .08 (.05) | (-.15, .33) | .48 |
| $a_i$ on Age | -.05 (-.01) | (-.16, .08) | .55 |
| $b_i$ on Age | -.09 (-.04) | (-.24, .02) | .12 |
| $c_i$ on Age | -.07 (-.06) | (-.17, .02) | .12 |
| $a_i$ on Sex | -.01 (-.01) | (-.16, .08) | .75 |
| $b_i$ on Sex | -.03 (-.03) | (-.14, .10) | .69 |
| $c_i$ on Sex | .04 (.08) | (-.07, .18) | .42 |
| <u>Within-Person</u> | | | |
| Share on Exchange ($a_i$) | -.05 (-.03) | (-.06, -.03) | < .001 |
| Share on Reputation ($b_i$) | -.08 (-.16) | (-.09, -.06) | < .001 |
| Share on Trustee Decision (*t-1*) ($c_i$) | .16 (.30) | (.13, .18) | < .001 |

*Note.* Except for the intercept of the decision to share, all standard deviations and credibility intervals refer to standardized regression coefficients. Italicized *a, b,* and *c* parameters represent random slopes included in the model. In MSEM, random slopes reflect between-person differences in the strength of a within-person effect (e.g., $b_i$ captures individual differences in the effect of reputation on decisions to share), which can then be related to other between-person predictors. Correspondence between random slopes and within-person associations is denoted in the parentheticals in the Within-Person section of the table. Within-person $n$ = 24192. Between-persons $n$ = 168.

**Section XI. Association between Personality and Prediction Error Modulation in the Default Network.**

Table S9. *Effect of Antagonism and its Facets on Prediction Error Modulation in the Default Network.*

| | Antagonism Model | | | | | | | |
|---|---|---|---|---|---|---|---|---|
| | Default Network | | Core | | Medial Temporal | | Dorsal Medial | |
| Variable | *B (SE)* | *p* | *B (SE)* | *p* | *B (SE)* | *p* | *B (SE)* | *p* |
| Intercept | .25 (.18) | .16 | .20 (.18) | .25 | .22 (.17) | .20 | .22 (.18) | .22 |
| Age | .10 (.08) | .26 | .07 (.08) | .43 | .09 (.08) | .25 | .10 (.08) | .22 |
| Sex | -.32 (.20) | .11 | -.28 (.20) | .16 | -.29 (.20) | .14 | -.27 (.20) | .18 |
| Antagonism | -.10 (.08) | .22 | -.08 (.08) | .31 | -.14 (.08) | .09 | -.04 (.08) | .66 |
| | Facets Model | | | | | | | |
| Intercept | .26 (.17) | .13 | .21 (.17) | .20 | .24 (.16) | .15 | .22 (.17) | .20 |
| Age | .18 (.08) | .03 | .15 (.19) | .07 | .19 (.08) | .02 | .16 (.08) | .06 |
| Sex | -.35 (.19) | .07 | -.31 (.19) | .10 | -.33 (.18) | .08 | -.28 (.20) | .15 |
| Callousness | -.39 (.10) | <.001 | -.40 (.10) | <.001 | -.46 (.09) | < .001 | -.25 (.10) | .01 |
| Exploit. | .26 (.10) | .01 | .28 (.10) | .01 | .29 (.10) | .003 | .20 (.10) | .05 |

Note. *n = 144* due to listwise-deletion of missing data. Age and traits were standardized prior to inclusion in the models.

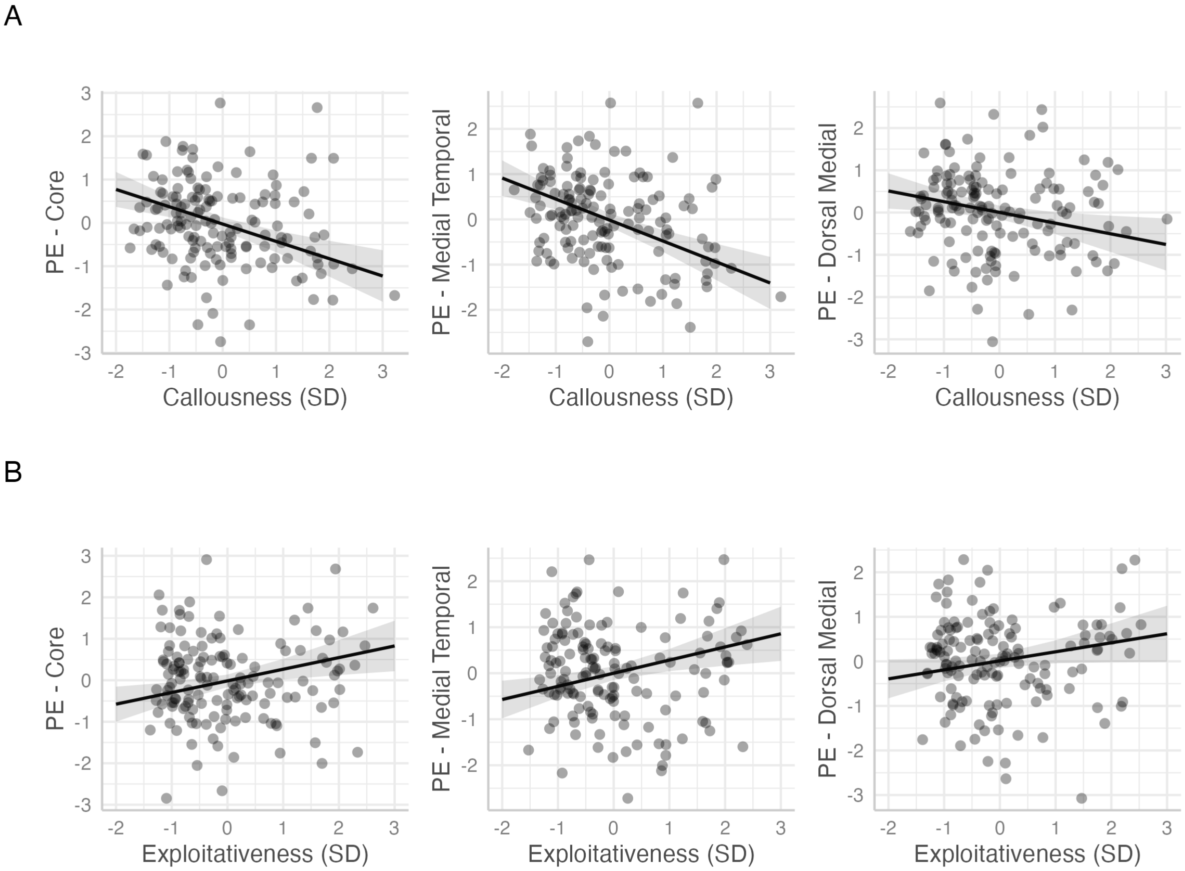

**Figure S6.** Association between A) callousness and B) exploitativeness and policy prediction errors in the core, medial temporal, and dorsal medial subsystems of the default network. All effects are significant at $p < .05$ controlling for age, sex, and the opposing facet, except the effect of exploitativeness on PEs in the dorsal medial subsystem, which is qualitatively the same ($\beta$ = .20, *SE* = .10, $p = .05$).

**Section XII. Effect of Personality on Behavior via Prediction Error Modulation of the Default Network**

The differential associations between the facets of antagonism and PE modulation in the default network implicate learning as a possible neural mechanism related both to self-reported personality and reciprocal decision-making during social exchanges. To formally test this, we extended our initial MSEM (Table S7), allowing the two facets of antagonism to predict PE modulation of the default network and testing whether prediction error signaling *mediated* the effect of callousness and/or exploitativeness on reciprocity. As in our previous MSEM, PEs encoded in the default network predicted a stronger effect of the trustee's last decision on the participant's subsequent decision to keep or share (a stronger reciprocity effect; $\beta$ = .25, 95% CI = .09, .42, $p$ < .001). Callousness was negatively ($\beta$ = -.36, 95% CI = -.50, -.20, $p$ < .001), and exploitativeness positively ($\beta$ = .24, 95% CI = .04, .39, $p$ = .03), associated with PE modulation in the default network (Table S10). Furthermore, there was a significant indirect effect of both callousness and exploitativeness on reciprocity via PE signaling (callousness: $b$ = -.05, 95% CI = -.10, -.02, $p$ < .001; exploitativeness: $b$ = .03, 95% CI = .004, .07, $p$ = .03).

Table S10. *Multilevel Structural Equation Model Examining Effect of Callousness and Exploitativeness on Behavior via Prediction Error Modulation of Default Network*

| Variable | *β (B)* | *95% CI* | *p* |
|---|---|---|---|
| Between-Person | | | |
| Share Intercept | --- (.06) | (-.11, .22) | .56 |
| Share on PE-Default Network | .11 (.06) | (-.07, .28) | .35 |
| Share on Callousness | -.07 (-.04) | (-.24, .11) | .50 |
| Share on Exploitativeness | -.12 (-.06) | (-.27, .08) | .29 |
| Share on Age | .21 (.17) | (.09, .32) | < .001 |
| Share on Sex | -.06 (-.11) | (-.16, .06) | .36 |
| $a_i$ on PE- Default Network | .01 (.00) | (-.21, .26) | .92 |
| $b_i$ on PE- Default Network | -.04 (-.01) | (-.25, .19) | .71 |
| $c_i$ on PE- Default Network | .25 (.13) | (.09, .42) | < .001 |
| $a_i$ on Callousness | .05 (.01) | (-.24, .26) | .67 |
| $b_i$ on Callousness | .21 (.07) | (-.05, .42) | .08 |
| $c_i$ on Callousness | -.03 (-.02) | (-.21, .15) | .75 |
| $a_i$ on Exploitativeness | -.04 (-.01) | (-.28, .18) | .72 |
| $b_i$ on Exploitativeness | .05 (.02) | (-.20, .27) | .65 |
| $c_i$ on Exploitativeness | -.16 (-.09) | (-.37, .05) | .14 |
| $a_i$ on Age | -.03 (-.01) | (-.19, .11) | .73 |

| | | | |
|---|---|---|---|
| $b_i$ on Age | -.09 (-.04) | (-.23, .05) | .18 |
| $c_i$ on Age | -.09 (-.07) | (-.20, .03) | .09 |
| $a_i$ on Sex | -.03 (-.01) | (-.15, .08) | .61 |
| $b_i$ on Sex | -.01 (-.01) | (-.15, .13) | .93 |
| $c_i$ on Sex | .04 (.07) | (-.08, .16) | .50 |
| PE-Default Network on Callousness | -.36 (-.38) | (-.50, -.20) | < .001 |
| PE-Default Network on Exploitativeness | .24 (.25) | (.04, .39) | .03 |
| PE-Default Network on Age | .11 (.17) | (-.01, .21) | .06 |
| PE-Default Network on Sex | -.10 (-.35) | (-.20, .01) | .06 |
| *Indirect Effects* | | | |
| Callousness → PE-DN → $c_i$ | --- (-.05) | (-.10, -.02) | < .001 |
| Exploitativeness → PE-DN → $c_i$ | --- (.03) | (.004, .07) | .03 |
| Within-Person | | | |
| Share on Exchange ($a_i$) | -.05 (-.02) | (-.06, -.03) | < .001 |
| Share on Reputation ($b_i$) | -.08 (-.17) | (-.09, -.05) | < .001 |
| Share on Trustee Decision (*t-1*) ($c_i$) | .16 (.33) | (.13, .18) | < .001 |

*Note.* Except for the intercept of the decision to share and indirect effects, all standard deviations and credibility intervals refer to standardized regression coefficients. Italicized *a, b,* and *c* parameters represent random slopes included in the model. In MSEM, random slopes reflect between-person differences in the strength of a within-person effect (e.g., $b_i$ captures individual differences in the effect of reputation on decisions to share), which can then be related to other between-person predictors. Correspondence between random slopes and within-person associations is denoted in the parentheticals in the Within-Person section of the table. Within-person $n$ = 24192. Between-persons $n$ = 168.

In previous analyses, we also found that prediction error signaling in the medial temporal subsystem was uniquely related to the effect of reputation on participant decisions to share (Table S8). Based on this finding, we examined whether prediction error signaling specifically in the medial temporal subsystem accounts for an association between antagonism and/or its facets and the effect of reputation on social decision-making.

As seen in Table S11 below, callousness and exploitativeness were differentially associated with prediction error signaling in the medial temporal subsystem negatively (callousness: $\beta$ = -.45, 95% CI = -.58, -.32, $p$ < .001; exploitativeness: ($\beta$ = .29, 95% CI = .13, .41, $p$ < .001), which in turn was marginally related to a stronger effect of reputation on decisions to share ($\beta$ = -.27, 95% CI = -.59, .01, $p$ = .06). Consequently, there was a marginal indirect effect such that PE signaling in the medial temporal subsystem mediated an association between callousness and reputation ($b$ = .04, 95% CI = -.002, .09, $p$ = .06). This is indicative of learning signals in the medial temporal subsystem dampening the effect of a good/neutral reputation on increased sharing among individuals high in callousness (note this indirect effect is positive because the good/neutral reputation is coded as 0, relative to the bad reputation, which was coded as 1). Prediction error signaling in the medial temporal subsystem also mediated an association between exploitativeness and the reputation effect, though once again, this effect was marginal ($b$ = -.03, 95% CI = -.07, .001, $p$ = .06). This indicates that higher exploitativeness was associated with a stronger effect of the good/neutral reputation on decisions to share, in part due to learning signals encoded in the medial temporal subsystem. Nonetheless, neither indirect effect was statistically significant at the a priori defined alpha of .05. At best, these results should be interpreted as tenuous evidence for prediction errors encoded in the medial temporal subsystem driving a link between facets of antagonism and the effect of reputation on social decision-making.

Table S11. *Multilevel Structural Equation Model Examining Effect of Callousness and Exploitativeness on Behavior via Prediction Error Modulation of Default Network Subsystems*

| Variable | *β (B)* | *95% CI* | *p* |
|---|---|---|---|
| Between-Person | | | |
| Share Intercept | -- (.06) | -.09, .25 | .52 |
| Share on PE-Core | .06 (.03) | (-.29, .34) | .74 |
| Share on PE-Medial Temporal | .12 (.06) | (-.12, .35) | .30 |
| Share on PE-Dorsal Medial | .00 (-.001) | (-.25, .23) | .99 |
| Share on Callousness | -.02 (-.01) | (-.20, .15) | .82 |
| Share on Exploitativeness | -.12 (-.07) | (-.33, .05) | .18 |
| Share on Age | .19 (.16) | (.08, .31) | < .001 |
| Share on Sex | -.05 (-.09) | (-.16, .05) | .30 |
| $a_i$ on PE-Core | .02 (.002) | (-.35, .34) | .88 |
| $b_i$ on PE-Core | -.02 (-.01) | (-.48, .37) | .92 |
| $c_i$ on PE-Core | .07 (.04) | (-.24, .38) | .76 |

| | | | |
|---|---|---|---|
| $a_i$ on PE-Medial Temporal | .16 (.02) | (-.17, .41) | .29 |
| $b_i$ on PE-Medial Temporal | -.27 (-.09) | (-.59, .01) | .06 |
| $c_i$ on PE-Medial Temporal | .19 (.10) | (-.09, .45) | .15 |
| $a_i$ on PE-Dorsal Medial | -.10 (-.01) | (-.38, .14) | .49 |
| $b_i$ on PE-Dorsal Medial | .12 (.04) | (-.18, .44) | .42 |
| $c_i$ on PE-Dorsal Medial | .08 (.04) | (-.24, .32) | .52 |
| $a_i$ on Callousness | .07 (.01) | (-.19, .28) | .59 |
| $b_i$ on Callousness | .09 (.03) | (-.21, .33) | .50 |
| $c_i$ on Callousness | .03 (.01) | (-.21, .22) | .85 |
| $a_i$ on Exploitativeness | -.09 (-.01) | (-.25, .17) | .48 |
| $b_i$ on Exploitativeness | .09 (.03) | (-.14, .32) | .54 |
| $c_i$ on Exploitativeness | -.19 (-.11) | (-.39, .002) | .06 |
| $a_i$ on Age | -.05 (-.01) | (-.17, .08) | .54 |
| $b_i$ on Age | -.09 (-.05) | (-.23, .04) | .20 |
| $c_i$ on Age | -.10 (-.08) | (-.22, .01) | .11 |
| $a_i$ on Sex | -.02 (-.01) | (-.14, .13) | .81 |
| $b_i$ on Sex | -.01 (-.01) | (-.14, .11) | .90 |
| $c_i$ on Sex | .04 (.08) | (-.06, .16) | .46 |
| PE-Core on Callousness | -.40 (-.43) | (-.52, -.22) | < .001 |
| PE- Core on Exploitativeness | .28 (.31) | (.12, .42) | < .001 |
| PE- Core on Age | .09 (.14) | (.001, .20) | .04 |
| PE- Core on Sex | -.08 (-.30) | (-.17, .02) | .08 |
| PE-Medial Temporal on Callousness | -.45 (-.49) | (-.58, -.32) | < .001 |
| PE-Medial Temporal on Exploitativeness | .29 (.31) | (.13, .41) | < .001 |
| PE-Medial Temporal on Age | .12 (.18) | (.02, .22) | .01 |
| PE-Medial Temporal on Sex | -.09 (-.32) | (-.18, .01) | .10 |
| PE-Dorsal Medial on Callousness | -.26 (-.26) | (-.43, -.10) | < .001 |
| PE- Dorsal Medial on Exploitativeness | .22 (.22) | (.05, .37) | < .001 |
| PE- Dorsal Medial on Age | .10 (.15) | (.01, .21) | .03 |
| PE- Dorsal Medial on Sex | -.09 (-.30) | (-.21, .03) | .16 |

| | | | |
|---|---|---|---|
| *Indirect Effects* | | | |
| Callousness → PE-MT → Reputation | .04 | (-.002, .09) | .06 |
| Exploitativeness → PE-MT → Reputation | -.03 | (-.07, .001) | .06 |
| Within-Person | | | |
| Share on Exchange ($a_i$) | -.05 (-.03) | (-.06, -.03) | < .001 |
| Share on Reputation ($b_i$) | -.08 (-.17) | (-.09, -.06) | < .001 |
| Share on Trustee Decision (*t-1*) ($c_i$) | .16 (.31) | (.14, .18) | < .001 |

*Note.* Except for the intercept of the decision to share, all standard deviations and credibility intervals refer to standardized regression coefficients. Italicized *a, b,* and *c* parameters represent random slopes included in the model. In MSEM, random slopes reflect between-person differences in the strength of a within-person effect (e.g., $b_i$ captures individual differences in the effect of reputation on decisions to share), which can then be related to other between-person predictors. Correspondence between random slopes and within-person associations is denoted in the parenthetical in the Within-Person section of the table. Within-person $n$ = 24192. Between-persons $n$ = 168.

**Section XIII: Direct Effects of Personality on Behavior**

To examine the effect of antagonism on behavioral performance, we fit a multilevel structural equation model (MSEM) in which individual differences in antagonism were allowed to predict trial-level variability (random slopes) in the effect of exchange, reputation, and trustee's previous decision on participant decisions to keep or share (coded 0/1, respectively; Table S12). Antagonism was associated with a weaker effect of the trustee's previous decision on the participant's decision to share (i.e., a weaker reciprocity effect) ($\beta$ = -.20, 95% CI = [-.37, -.04], *p* = .01; Figure S7A). More specifically, the trustee's decision to share on the previous trial was associated with a higher likelihood of the participant reciprocating, but only for participants who scored below the 88$^{th}$ percentile on antagonism ($z$ < 1.17). Antagonism also moderated the effect of reputation on sharing ($\beta$ = .22, 95% CI = [.02, .37], *p* = .02; see Fig S7B), such that participants who were low (-1 *SD*; *B* = -.23, 95% CI = [-.39, -.11], *p* < .001), or average (*B* = -.17, 95% CI = [-.30, -.06], *p* < .001) on antagonism were less likely to share with the bad trustee relative to the good and neutral trustees, whereas reputation had no effect on sharing for those high (+1 *SD*) in antagonism (*B* = -.10, 95% CI = [-.22, .02], *p* = .14).

A second MSEM was conducted to examine the effect of exploitativeness and callousness on behavioral performance in the trust game (Table S12). When both facets were entered into the model as simultaneous predictors, callousness, but not exploitativeness, significantly moderated the effect of reputation on sharing ($\beta$ = .26, 95% CI = [.07, .47], *p* = .01; Fig S7C). Participants who were low (-1 *SD*; *B* = -.29, 95% CI = [-.43, -.14], p = < .001) or average (*B* = -.20, 95% CI = [-.31, -.08], p = < .001) on callousness were less likely to share with the bad trustee relative to the good and neutral trustees, whereas reputation had no effect on sharing for those high (+1 *SD*) in callousness (*B* = -.12, 95% CI = [-.24, .004], *p* = .07). Neither facet significantly moderated the effect of reciprocity on participants' decision to share. When entered independently, the effect of each facet on reciprocity was similar to that of antagonism (Table S13). Taken together, these facet-level results suggest that the moderating effect of antagonism on reputation is primarily driven by its callousness facet, whereas the moderating effect of antagonism on reciprocity is not better accounted for by either of its unique facets.

Table S12. *Multilevel Structural Equation Model Examining Moderation of the Effects of Reputation and Reciprocity on Decision to Share by Antagonism and its Facets*

| | Antagonism Model | | | Facets Model | | |
|---|---|---|---|---|---|---|
| Variable | *β (B)* | *95% CI* | *p* | *β (B)* | *95% CI* | *p* |
| Between-Person | | | | | | |
| Share Intercept | -- (.06) | -.08, .24 | .42 | -- (.07) | -.08, .23 | .40 |
| Share on Antagonism | -.17 (-.09) | -.30, .001 | .06 | --- | --- | --- |
| Share on Callousness | --- | --- | --- | -.11 (-.06) | -.28, .08 | .23 |
| Share on Exploitativeness | --- | --- | --- | -.09 (-.05) | -.27, .10 | .43 |
| Share on Age | .21 (.17) | .11, .31 | <.001 | .22 (.18) | .11, .33 | <.001 |
| Share on Sex | -.06 (-.11) | -.18, .05 | .25 | -.07 (-.13) | -.16, .03 | .19 |

| | | | | | | |
|---|---|---|---|---|---|---|
| $a_i$ on Antagonism | -.03 (-.00) | -.19, .17 | .80 | --- | --- | --- |
| $b_i$ on Antagonism | .22 (.07) | .02, .37 | .02 | --- | --- | --- |
| $c_i$ on Antagonism | -.20 (-.11) | -.37, -.04 | .01 | --- | --- | --- |
| $a_i$ on Callousness | --- | --- | --- | .07 (.01) | -.14, .28 | .49 |
| $b_i$ on Callousness | --- | --- | --- | .26 (.09) | .04, .45 | .01 |
| $c_i$ on Callousness | --- | --- | --- | -.13 (-.07) | -.32, .07 | .20 |
| $a_i$ on Exploitativeness | --- | --- | --- | -.06 (-.01) | -.28, .19 | .58 |
| $b_i$ on Exploitativeness | --- | --- | --- | -.01 (-.00) | -.24, .22 | .91 |
| $c_i$ on Exploitativeness | --- | --- | --- | -.11 (-.06) | -.28, .08 | .33 |
| $a_i$ on Age | -.05 (-.01) | -.17, .09 | .51 | -.05 (-.01) | -.20, .10 | .49 |
| $b_i$ on Age | -.09 (-.04) | -.25, .05 | .24 | -.14 (-.06) | -.25, -.02 | .02 |
| $c_i$ on Age | -.08 (-.06) | -.19, .02 | .12 | -.07 (-.05) | -.19, .05 | .23 |
| $a_i$ on Sex | -.03 (-.01) | -.16, .08 | .62 | -.02 (-.01) | -.15, .11 | .78 |
| $b_i$ on Sex | -.00 (-.00) | -.13, .13 | .99 | .01 (.01) | -.12, .14 | .89 |
| $c_i$ on Sex | .02 (.04) | -.09, .13 | .64 | .01 (.02) | -.10, .12 | .82 |
| Within-Person | | | | | | |
| Share on Exchange ($a_i$) | -.05 (-.03) | -.07, -.03 | <.001 | -.05 (-.03) | -.06, -.03 | <.001 |
| Share on Reputation ($b_i$) | -.08 (-.17) | -.10, -.06 | <.001 | -.08 (-.19) | -.10, -.06 | <.001 |
| Share on Trustee Decision ($t-1$) ($c_i$) | .16 (.34) | .13, .18 | <.001 | .16 (.36) | .14, .18 | <.001 |

*Note.* Except for the intercept of the decision to share, all standard deviations and credibility intervals refer to standardized regression coefficients. Italicized *a, b,* and *c* parameters represent random slopes included in the model. In MSEM, random slopes reflect between-person differences in the strength of a within-person effect (e.g., $b_i$ captures individual differences in the effect of reputation on decisions to share), which can then be related to other between-person predictors. Correspondence between random slopes and within-person associations is denoted in the parentheticals in the Within-Person section of the table. Within-person $n$ = 24192. Between-persons $n$ = 168.

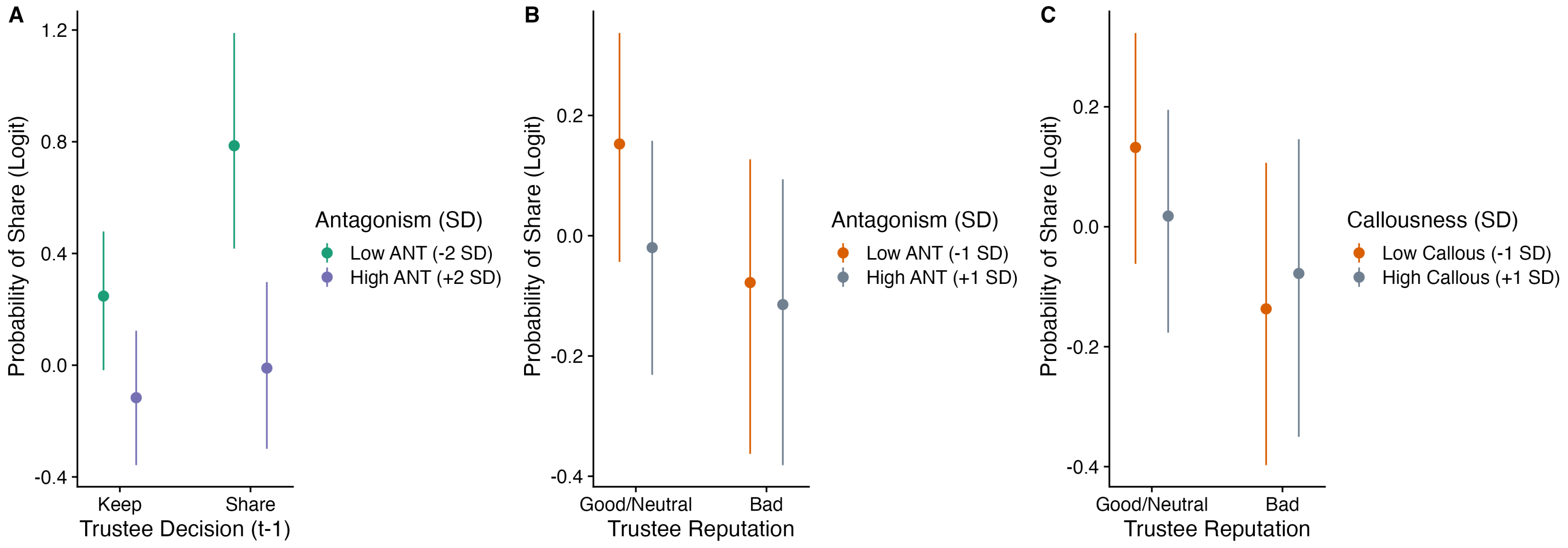

**Figure S7.** Trait moderation of behavioral effects on the modified social trust task.

Table S13. *Multilevel Structural Equation Model Examining Moderation of the Effects of Reputation and Reciprocity on Decision to Share by Individual Antagonism Facets*

| | Callousness Model | | | Exploitativeness Model | | |
|---|---|---|---|---|---|---|
| Variable | *β (B)* | *95% CI* | *p* | *β (B)* | *95% CI* | *p* |
| Between-Person | | | | | | |
| Share Intercept | --- (.07) | -.08, .24 | .36 | --- (.06) | -.09, .23 | .48 |
| Share on Facet | -.15 (-.08) | -.29, .01 | .09 | -.15 (-.08) | -.29, .03 | .11 |
| Share on Age | .23 (.19) | .13, .32 | <.001 | .21 (.16) | .11, .31 | <.001 |
| Share on Sex | -.06 (-.12) | -.18, .04 | .21 | -.06 (-.11) | -.18, .05 | .28 |
| $a_i$ on Facet | .002 (.00) | -.17, .19 | .98 | -.03 (-.00) | -.20, .14 | .82 |
| $b_i$ on Facet | .25 (.08) | .06, .40 | .01 | .14 (.04) | -.06, .30 | .19 |
| $c_i$ on Facet | -.18 (-.10) | -.36, -.03 | .02 | -.17 (-.09) | -.35, -.02 | .03 |
| $a_i$ on Age | -.05 (-.01) | -.17, .10 | .54 | -.05 (-.01) | -.18, .09 | .50 |
| $b_i$ on Age | -.12 (-.05) | -.27, .01 | .07 | -.09 (-.04) | -.25, .05 | .23 |
| $c_i$ on Age | -.06 (-.04) | -.17, .04 | .35 | -.09 (-.07) | -.20, .02 | .12 |
| $a_i$ on Sex | -.03 (-.01) | -.16, .08 | .67 | -.03 (-.01) | -.16, .08 | .62 |
| $b_i$ on Sex | .01 (.01) | -.11, .14 | .91 | -.01 (-.01) | -.13, .12 | .88 |
| $c_i$ on Sex | .02 (.03) | -.10, .13 | .72 | .03 (.05) | -.08, .13 | .54 |
| Within-Person | | | | | | |
| Share on Exchange ($a_i$) | -.05 (-.03) | -.07, -.03 | <.001 | -.05 (-.03) | -.07, -.03 | <.001 |
| Share on Reputation ($b_i$) | -.08 (-.17) | -.10, -.06 | <.001 | -.08 (-.16) | -.10, -.06 | <.001 |
| Share on Trustee Decision (*t-1*) ($c_i$) | .16 (.34) | .13, .18 | <.001 | .16 (.33) | .13, .18 | <.001 |

*Note.* Except for the intercept of the decision to share, all standard deviations and credibility intervals refer to standardized regression coefficients. Italicized *a, b,* and *c* parameters represent random slopes included in the model. In MSEM, random slopes reflect between-person differences in the strength of a within-person effect (e.g., $b_i$ captures individual differences in the effect of reputation on decisions to share), which can then be related to other between-person predictors. Correspondence between random slopes and within-person associations is denoted in the parentheticals in the Within-Person section of the table. Within-person $n$ = 24192. Between-persons $n$ = 168.

**Section XIV. Effect of Antagonism on Behavior via Prediction Errors in the Default Network**

Our primary interest was in understanding whether callousness and exploitativeness were differentially related to learning, and in turn, decision-making during a social exchange. As a secondary analysis, we also examined links between broadband antagonism, learning on the task, and trial-level decision-making, Specifically, we allowed individual differences in antagonism to predict prediction error modulation of the default network, which in turn moderated trial-level effects of exchange number, reputation, and reciprocity. As noted in our main findings, PEs encoded in the default network predicted a stronger effect of the trustee's last decision on the participant's subsequent decision to keep or share (a stronger reciprocity effect; $\beta$ = .24, 95% CI = .05, .38, $p$ = .01). Separately, higher antagonism was associated with a weaker effect of the trustee's previous decision (i.e., a weaker reciprocity effect) on the participant's decision to share ($\beta$ = -.17, 95% CI = -.32, -.02, $p$ = .03). However, there was no association between antagonism and prediction error signaling in the overall default network. Consistent with this, learning signals encoded in the default network did not appear to account for the association between antagonism and the reciprocity effect on the task (indirect effect of antagonism to reciprocity via PE-DN: $B$ = -.01, $p$ = .21; see Table S12 below).

Table S14. *MSEM Examining Effect of Antagonism on Behavior via PE Modulation of the Default Network*

| Variable | *β (B)* | 95% CI | *p* |
|---|---|---|---|
| Between-Person | | | |
| Share Intercept | --- (.06) | -.10, .26 | .62 |
| Share on PE-DN | .08 (.05) | -.07, .23 | .21 |
| Share on Antagonism | -.14 (-.08) | -.30, .00 | .07 |
| Share on Age | .22 (.17) | .10, .32 | < .001 |
| Share on Sex | -.06 (-.11) | -.17, .07 | .31 |
| $a_i$ on PE-DN | .00 (.00) | -.19, .17 | .99 |
| $a_i$ on Antagonism | -.03 (-.00) | -.22, .16 | .80 |
| $a_i$ on Age | -.04 (-.01) | -.18, .09 | .55 |
| $a_i$ on Sex | -.03 (-.01) | -.15, .09 | .65 |
| $b_i$ on PE-DN | -.09 (-.03) | -.28, .09 | .29 |
| $b_i$ on Antagonism | .18 (.06) | -.01, .37 | .06 |
| $b_i$ on Age | -.10 (-.04) | -.22, .05 | .17 |
| $b_i$ on Sex_1 | -.01 (-.01) | -.14, .12 | .90 |
| $c_i$ on PE-DN | .24 (.13) | .05, .38 | .01 |
| $c_i$ on Antagonism | -.17 (-.09) | -.32, -.02 | .03 |

| | | | |
|---|---|---|---|
| $c_i$ on Age | -.08 (-.06) | -.19, .01 | .11 |
| $c_i$ on Sex | .04 (.08) | -.06, .16 | .39 |
| PE-DN on Antagonism | -.10 (-.10) | -.25, .06 | .20 |
| PE-DN on Age | .06 (.09) | -.06, .16 | .34 |
| PE-DN on Sex | -.10 (-.33) | -.22, -.01 | .04 |
| Indirect Effects | | | |
| Antagonism → PE-DN → $b_i$ | --- (.00) | -.004, .01 | .47 |
| Antagonism → PE-DN → $c_i$ | --- (-.01) | -.04, .01 | .21 |
| Within-Person | | | |
| Share on Exchange ($a_i$) | -.05 (-.03) | -.07, -.03 | < .001 |
| Share on Reputation ($b_i$) | -.08 (-.16) | -.10, -.06 | < .001 |
| Share on Trustee Decision (*t-1*) ($c_i$) | .16 (.32) | .13, .18 | < .001 |

*Note.* Except for the intercept of the decision to share and indirect effects, all standard deviations and credibility intervals refer to standardized regression coefficients. Italicized *a, b,* and *c* parameters represent random slopes included in the model. In MSEM, random slopes reflect between-person differences in the strength of a within-person effect (e.g., $b_i$ captures individual differences in the effect of reputation on decisions to share), which can then be related to other between-person predictors. Correspondence between random slopes and within-person associations is denoted in the parentheticals in the Within-Person section of the table. Within-person $n$ = 24192. Between-persons $n$ = 168.
